# Comprehensive analysis of TEAD inhibition in meningioma identifies MEK and mTOR inhibition as effective combination therapies against resistant lines

**DOI:** 10.64898/2026.03.20.713271

**Authors:** Dylan J Keiser, Meghan S Buddy, Derek J Sauer, Solmaz Mojarad-Jabali, Elena Ibanez, Qing Li, Daniel Rodriguez-Vazquez, Missia Kohler-Skinner, Abigail G Parrish, Mona Foth, David Gillespie, Arvonn Tully, Renee D Read, David Nix, Sheri Holmen, Howard Colman, William Couldwell, Randy L Jensen, Martin Distel, Frank Szulzewsky

## Abstract

Meningiomas are the most common primary central nervous system tumors in adults, posing a significant burden to society. Although a large percentage of lower-grade meningiomas are curable by surgery or radiation alone, high-grade and a subset of low-grade meningiomas demonstrate recurrences and complications from treatment. Systemic therapies for meningioma remain ineffective, and no targeted treatments are approved. We have recently shown that YAP1/TAZ-TEAD signaling is an oncogenic driver of meningioma. Despite the central role of YAP1/TAZ-TEAD signaling in NF2-deficient/mutant tumors, including meningioma, no studies have systematically examined TEAD inhibition across molecularly defined meningioma subtypes or investigated mechanisms of resistance in this disease. Here, using established and patient-derived meningioma cell lines, we demonstrate that genetic ablation of YAP1/TAZ suppresses growth in both NF2 mutant and NF2 wild type cell lines, establishing YAP1/TAZ-TEAD signaling as a shared oncogenic dependency. Pharmacologic TEAD inhibition suppressed growth of NF2 mutant and NF2 wild type meningioma cell lines alike, while higher levels of genomic rearrangement and mutations in the MAPK pathway contributed to TEADi resistance. RNA-Seq and Western Blot analysis identified compensatory activation of MEK-ERK, mTOR-S6, and FAK signaling in resistant lines. Importantly, co-targeting these pathways in combination with TEADi was able to overcome TEADi resistance and was superior to MEK/mTOR/FAK inhibition alone. These studies provide a compelling proof-of-concept that TEADi represents a novel therapeutic vulnerability in meningioma and reveal adaptive signaling responses that can be therapeutically exploited.

## Introduction

Meningiomas are the most common primary central nervous system (CNS) tumors in adults, with an incidence of 9.73 per 100,000 people per year, comprising 37.6% of all primary CNS tumors ^1,2^. Incidence increases with age, with an incidence rate of 18.69 per 100,000 people per year in patients aged 40 and over ^3^. Despite being the most common primary CNS tumor in adults, meningiomas have been significantly understudied compared to other CNS malignancies, such as glioblastoma. The majority of meningiomas are generally benign and curable by surgery alone, however, around 15-20% are of higher-grade (WHO grades II and III), with a rising incidence ^4,5^. Higher-grade meningiomas frequently show aggressive and invasive growth, recur even after multiple rounds of surgery and radiation, and are ultimately fatal, highlighting the need for additional therapeutic options ^6^. In addition, a significant number of meningiomas are often less accessible for surgery due to their location and/or involvement of major blood vessels and cranial nerves, such as skull base or optic nerve sheath meningiomas, that limit complete resection and thereby lead to progression and worsening symptoms ^7^.

Chemotherapeutic agents used for the treatment of other solid tumors, including cyclophosphamide, doxorubicin, temozolomide, and vincristine, have been largely ineffective in meningiomas ^8–12^. The advance of high-throughput sequencing and large-scale genomic and epigenomic studies have led to the identification of clinically-relevant subtypes and molecular drivers of meningioma ^13,14^, enabling novel molecular classification and diagnostic approaches, reshaping the molecular taxonomy of meningioma. However, this has not yet been translated into effective targeted therapies. Targeted inhibitors, including FAK, PI3K, CDK4/6, and EZH2 inhibitors are currently undergoing clinical trials and may show efficacy in a subset of patients ^15–17^. Therefore, additional chemotherapeutic approaches are urgently needed for the treatment of meningiomas, especially for management of hard-to-treat high-grade meningioma subtypes that are either inaccessible to surgery or that recur even after multiple rounds of surgery and radiotherapy.

Recurrent somatic alterations in meningiomas include mutations in NF2, TRAF7, KLF4, AKT1, SMO, POLR2A, and BAP1, as well as chromosomal copy number changes and DNA methylation-based subgroups that more accurately predict clinical outcome than histology alone ^13,14,18–21^. Among these alterations, inactivation of the NF2 gene is the most frequent event, occurring in approximately 40–60% of sporadic meningiomas ^14,19,22^. Furthermore, patients with the neurofibromatosis type 2 syndrome, which is frequently caused by germline NF2 haploinsufficiency, develop multiple meningiomas ^23^. Importantly, NF2 loss converges functionally on dysregulation of the Hippo signaling pathway and activation of the transcriptional co-activators YAP1 (Yes-associated protein 1) and TAZ (encoded by the WWTR1 gene) ^24–26^, positioning the YAP1/TAZ-TEAD axis as a central driver in a substantial subset of meningiomas.

The Hippo pathway is an evolutionarily conserved kinase cascade that regulates organ size, tissue homeostasis, regeneration, and stem cell behavior ^27^. In its canonical configuration, upstream regulators including NF2 (Merlin), SAV1, and other polarity and cytoskeletal proteins promote activation of the MST1/2 kinases, which in turn phosphorylate and activate LATS1/2. Activated LATS kinases phosphorylate YAP1 and TAZ, resulting in their cytoplasmic retention, ubiquitin-mediated degradation, or functional inactivation ^27^. When Hippo signaling is suppressed, such as through genetic loss of NF2, YAP1 and TAZ accumulate in the nucleus. There, they bind members of the TEAD (TEA domain) family of transcription factors (TEAD1-4), which serve as the primary DNA-binding partners of YAP1/TAZ, and drive expression of genes involved in proliferation (e.g., CTGF, CYR61), survival, extracellular matrix remodeling, and stemness ^28^. In many solid tumors, persistent YAP1/TAZ-TEAD activity promotes oncogenic transformation, tumor growth, metastasis, and therapeutic resistance.

We and others have detected increased YAP1/TAZ-TEAD activity (measured via the expression of YAP1/TAZ-TEAD target genes, such as CTGF and CYR61) in NF2 mutant (NF2mut) meningioma tumor tissues and/or cell lines compared to NF2 wild type (NF2wt) meningiomas ^29–32^. Furthermore, the expression of a non-regulatable activated YAP1 variant (S127/397A-(2SA)-YAP1) induces transcriptional changes similar to functional NF2 loss in human neural stem cells and, in addition, the intracranial expression of 2SA-YAP1 in meningeal cells is sufficient to induce the formation of meningioma-like tumors in mice ^32^. Lastly, recurrent YAP1 gene fusions (most commonly YAP1-MAML2) have been identified in a subset of pediatric NF2wt meningiomas ^33^. The oncogenic functions of these YAP1 fusions fundamentally rely on their ability to exert deregulated YAP activity, and YAP1 fusion-positive human meningiomas resemble NF2mut meningiomas by both gene expression and DNA methylation-based classification ^32–34^. These data suggest that the functional NF2 loss in meningiomas induces oncogenic YAP1/TAZ-TEAD activity which may be a therapeutic target in a subset of these tumors. However, while benign (low-grade) NF2mut meningiomas harbor few mutations in addition to functional NF2 loss, aggressive (high-grade) NF2mut meningiomas frequently harbor additional mutations and copy number losses ^14,35,36^. The presence of these additional mutations may render aggressive-type NF2mut meningiomas less reliant on YAP1/TAZ-TEAD signaling. Consistent with this, we recently detected decreased levels of YAP1 activity (through the expression of direct YAP1 targets (e.g. CTGF)) in both bulk RNA-Seq samples as well as single-cell RNA-Seq samples of aggressive NF2mut meningiomas ^30^.

Given the prominent role of YAP1/TAZ-TEAD signaling in multiple cancers, therapeutic strategies targeting TEADs, posttranslational modifications of TEADs, or the interaction between TEADs and YAP1/TAZ have been developed ^37^. Structural and functional insights into TEAD biology have revealed druggable vulnerabilities, and current TEAD inhibitors generally either sterically block the interaction between TEAD and YAP1/TAZ (e.g. by occupying the TEAD Ω loop region) or by occupying the central hydrophobic pocket of TEAD, blocking TEAD auto-palmitoylation ^38–41^. Several pan-TEAD inhibitors (TEADi) have entered early-phase clinical trials (NCT04665206, NCT04857372) ^42^, particularly in tumors with NF2 loss or YAP1 or WWTR1/TAZ gene fusions (e.g. Epithelioid Hemangioendotheliomas). However, their clinical development faces important challenges, including the identification of robust predictive biomarkers of TEAD dependency, potential on-target toxicities given the role of YAP1/TAZ in normal tissue homeostasis and regeneration, and emergence of adaptive resistance mechanisms or pathway bypass signaling. Despite these challenges, these agents provide proof-of-concept that TEAD is pharmacologically tractable and targeting YAP1/TAZ-TEAD signaling in specific tumor populations may have clinical benefit and raise the possibility of extending this strategy to molecularly selected meningioma patients.

Despite the prominent role of YAP1/TAZ-TEAD signaling in meningioma, the efficacy of TEADi has not been extensively studied in meningioma. Furthermore, pathways that may contribute to resistance to TEADi therapy need to be explored in order to inform the development of effective combination approaches. Here, we explore the importance of YAP1/TAZ-TEAD signaling and the efficacy of TEADi, as well as potential combination therapies in multiple established and primary meningioma cell lines. First, we show that combined YAP1 and TAZ knockout results in highly significant growth reduction in all tested meningioma lines, including both NF2mut and NF2wt lines, suggesting that YAP1/TAZ-TEAD signaling is a central driver in meningiomas regardless of NF2 status. Second, in vitro treatment of 26 meningioma lines with four different TEAD inhibitors (IAG-933, MYF-03-176, GNE-7883, K-975) identified TEADi-sensitive and -resistant cell lines. While low-grade NF2mut and NF2wt meningioma cell lines were similarly sensitive to TEADi, increased levels of genomic rearrangement and mutations in MAPK pathway contributed to TEADi resistance. RNA-Seq and Western Blot analysis of both TEADi-sensitive and -resistant lines identified upregulation of MAPK and mTOR-S6 pathways in TEADi-resistant, but not -sensitive lines. The combination treatment of TEADi and mTORi or MEKi was able to overcome resistance to TEADi and yielded superior efficiency to either therapy alone. Finally, whole exome sequencing identified mutations in the MAPK and PI3K pathways in TEADi resistant lines. In summary, we establish YAP1/TAZ-TEAD signaling as a central driver in both NF2mut and NF2wt meningiomas and identify possible combination therapies that can overcome the resistance to TEADi.

## Results

### YAP1/TAZ expression is necessary for the growth of both NF2mut and NF2wt meningioma cells

To assess the importance of YAP1/TAZ-signaling for the growth of meningiomas, we used CRISPR-Cas9 gene editing to assess the effect of genetic ablation of YAP1/TAZ on the growth of meningioma cells. We tested the effects of combined YAP1 and TAZ double gene knockout compared to that of cells treated with CD8A control guides in multiple meningioma cell lines, including three NF2mut (one low-grade (BenMen1) and two high-grade lines (KT21-MG1, CH-157MN)) and one NF2wt line (IOMM-Lee). Successful gene knockout was confirmed by PCR/sanger sequencing (Suppl. Figure S1A). We observed a significant decrease in YAP1/TAZ-TEAD signaling, measured via the expression of the YAP1/TAZ-TEAD target genes CTGF, CYR61, ANKRD1, and AMOTL2 (Figure 1A; Suppl. Figure S1B). We performed live-cell imaging to analyze the impact of YAP1/TAZ double knockout on cell growth. We observed an almost complete growth inhibition upon YAP1 and TAZ double knockout in two of the three NF2mut cell lines (BenMen1, KT21-MG1). The cell confluence of BenMen1 and KT21-MG1 cells harboring a YAP1+TAZ double knockout did not significantly increase compared to t = 0 (1.075±0.08-fold increase (padj = 1) and 1.32±0.41-fold increase (padj = 0.98), respectively) (Figure 1B). We also observed a highly significant growth reduction in CH-157MN and IOMM-Lee cells upon co-deletion of YAP1 and TAZ (Figure 1B). However, both CH-157MN (1.37±0.04-fold increase (padj < 0.0001) and IOMM-Lee (5.72±1.56-fold increase (padj < 0.0001)) cell lines were able to significantly grow even upon YAP1 and TAZ double knockout. These results suggest that YAP1/TAZ-TEAD transcriptional activity is involved in the growth of meningioma cells (including high-grade NF2mut and NF2wt subtypes) and, therefore, is an attractive therapeutic target. However, while a subset of cells stop growing almost completely upon loss of YAP1/TAZ-TEAD signaling, other cell lines are able to partially overcome this effect.

**Figure 1:**
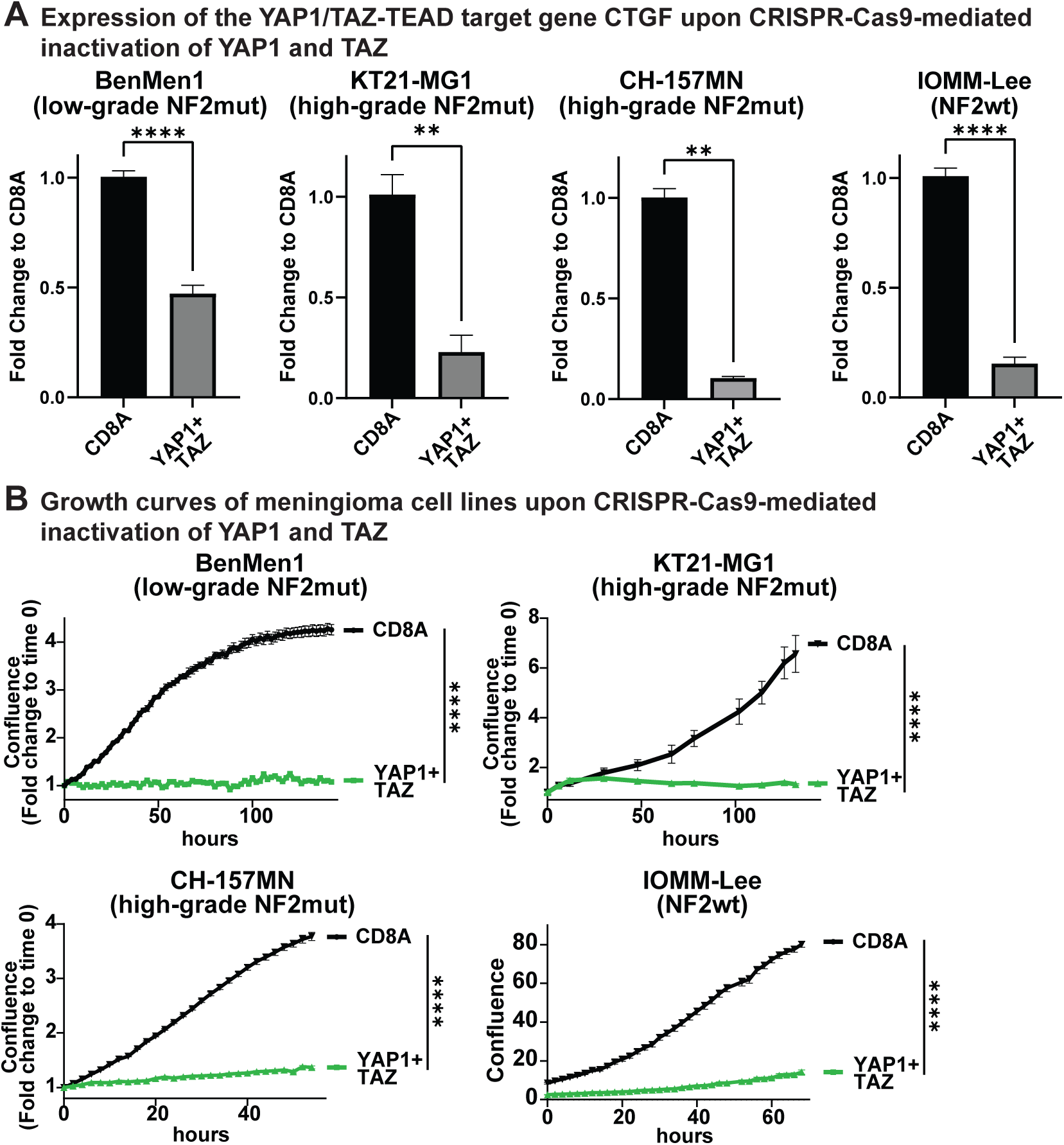
Combined loss of YAP1 and TAZ (WWTR1) significantly reduces the growth of both NF2 mutant and NF2 wild type meningioma cells. A) Expression of the YAP1/TAZ-TEAD target gene CTGF in benign NF2mut (BenMen1 (n = 10 each)), aggressive high-grade NF2mut (KT21-MG1 (n = 3 each), CH-157MN (n = 3 each)), and NF2wt (IOMM-Lee (n = 14 each)) meningioma cell lines upon CRISPR-Cas9-mediated inactivation of either CD8A (control) or combined inactivation of YAP1 and TAZ (WWTR1). B) Cell growth measured by live-cell imaging of benign NF2mut (BenMen1 (n = 5 each)), aggressive high-grade NF2mut (KT21-MG1 (n = 5 each), CH-157MN (n = 4 each)), and NF2wt (IOMM-Lee (n = 4 each)) meningioma cell lines upon CRISPR-Cas9-mediated inactivation of either CD8A (control) or combined inactivation of YAP1 and TAZ (WWTR1). Error bars show SEM. Analysis was done using a two-tailed t test (A) or ordinary two-way ANOVA (B, for the last time point).

### TEADi shows significant efficacy against benign and a subset of high-grade NF2 mutant meningioma cell lines

We collected 26 human meningioma cell lines, including four established lines (Ben-Men-1, KT21-MG1, CH-157MN, IOMM-Lee) and 22 primary lines, including both NF2mut and NF2wt cell lines. NF2/Merlin status was assessed via Western Blot staining (Suppl. Figure S2A). We then assessed the efficacy of four TEAD inhibitors (MYF-03-176, IAG-933, K-975, GNE-7883; dosing range of 1, 10, 50, 100, 200, 500, 1000, 2000, and 5000 nM, 0.1% DMSO as control)^38–41^ via live-cell imaging and CellTiter-Glo viability assays and determined absolute IC_50_ values for each inhibitor and cell line (n = 3 per cell line and inhibitor; Figure 2A-C; Suppl. Figure S2B-F). Cells were cultured until DMSO controls reached confluency of >80 percent. A specific cell line was deemed sensitive to a specific inhibitor if the absolute IC_50_ for a specific TEAD inhibitor was below 250 nM (determined by Cell Titer-Glo). We observed both sensitive and resistant NF2mut and NF2wt cell lines, suggesting that NF2 status is not a general indicator of TEADi sensitivity (Table 1). In general, IAG-933 and MYF-03-176 showed the highest efficacy in most cell lines (Figure 2A-C; Suppl. Figure S2B-F). Absolute IC_50_ values for IAG-933 ranged from 39-1500 nM, whereas absolute IC_50_ values for MYF-03-176 ranged from 76 nM to >5000 nM. In TEADi-sensitive lines both IAG-933 and MYF-03-176 showed somewhat comparable efficacies, whereas GNE-7883 and K-975 showed lower efficacy but still led to a significant growth reduction. By contrast, in TEADi-resistant lines all inhibitors displayed significantly lower efficacies (Figure 2A). However, while IAG-933, and to a lesser degree MYF-03-176, were still able to induce a significant growth reduction at higher concentrations, GNE-7883 and K-975 were frequently unable to exert any effect on the growth of resistant lines.

**Figure 2:**
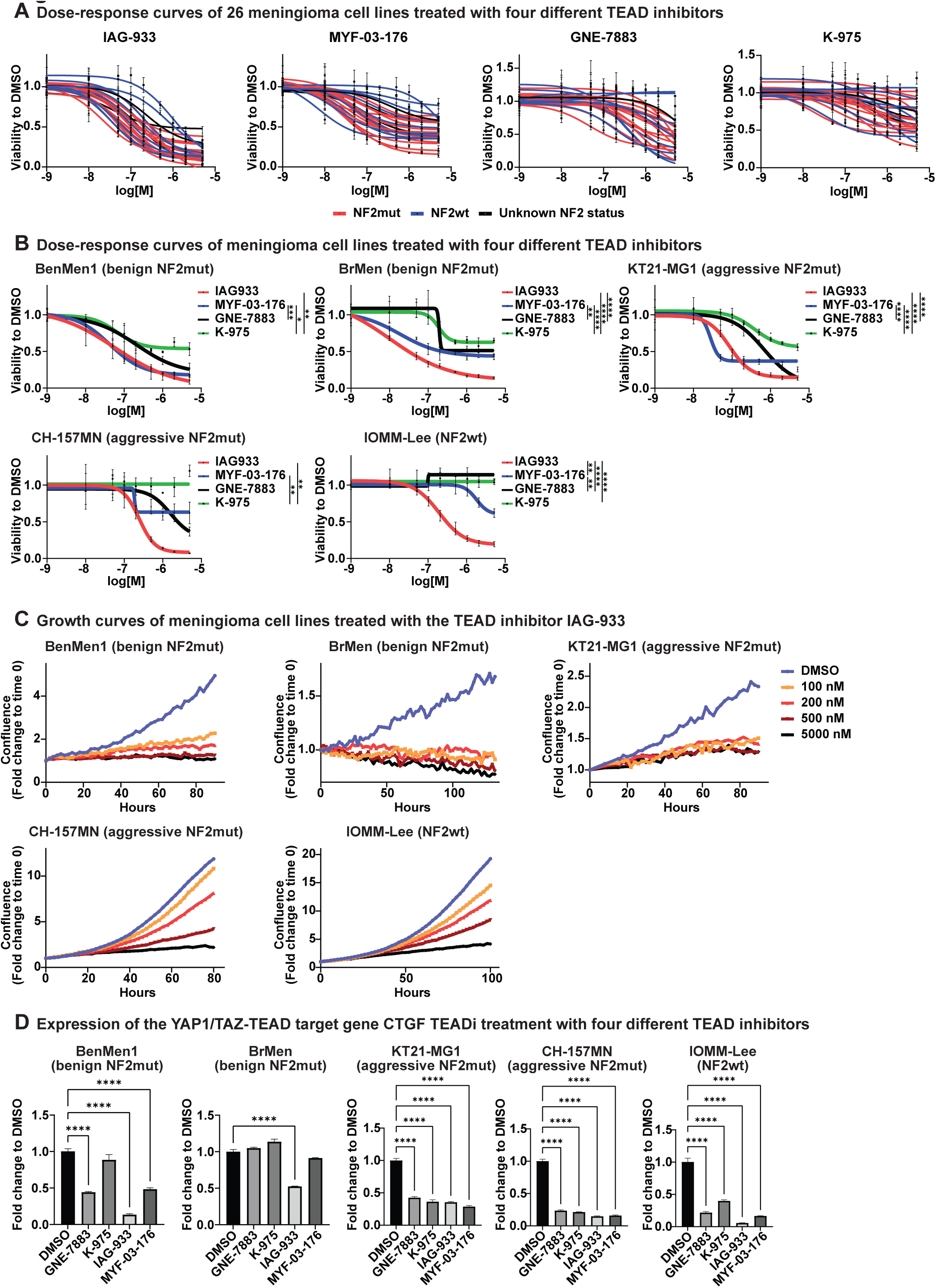
TEADi shows significant efficacy against benign and a subset of high-grade NF2 mutant meningioma cell lines. A) Cell viability dose response curves of 26 meningioma cell lines treated with different doses of four TEAD inhibitors (IAG-933, MYF-03-176, GNE-7883, K-975). Cell viability was measured upon the DMSO control reaching a confluence of > 80%. All experiments were performed in triplicates. B) Viability dose response curves of benign NF2mut (BenMen1, BrMen), aggressive high-grade NF2mut (KT21-MG1, CH-157MN), and NF2wt (IOMM-Lee) meningioma cell lines treated with different doses of four TEAD inhibitors (IAG-933, MYF-03-176, GNE-7883, K-975). All experiments were performed in triplicates. C) Growth curves measured by live-cell imaging of benign NF2mut (BenMen1, BrMen), aggressive high-grade NF2mut (KT21-MG1, CH-157MN), and NF2wt (IOMM-Lee) meningioma cell lines treated with different doses of the TEAD inhibitor IAG-933 (100, 200, 500, 1000 nM, 0.05% DMSO). All experiments were performed in triplicates. D) Expression of the YAP1/TAZ-TEAD target gene CTGF in benign NF2mut (BenMen1, BrMen), aggressive high-grade NF2mut (KT21-MG1, CH-157MN), and NF2wt (IOMM-Lee) meningioma cell lines upon treatment with four TEAD inhibitors (IAG-933 (500 nM), MYF-03-176 (500 nM), GNE-7883 (5000 nM), K-975 (1000 nM)). All experiments were performed in triplicates. Error bars show SD (A, B) or SEM (D). Analysis was done using an ordinary one-way ANOVA (D) or ordinary two-way ANOVA (A, B, comparison at 200 nM) with multiple comparisons testing.

**Table 1.** Overview of different meningioma cell lines tested (absolute IC50 values, NF2 status, WHO grade).

|  | IAG-933 | MYF-03-176 | GNE-7883 | K-975 | NF2 status | WHO grade |
|---|---|---|---|---|---|---|
| **BenMen1** | 78 | 76 | 410 | 2443 | NF2 MUT | WHO I |
| **BrMen** | 39 | 456 | 1629 | >5000 | NF2 MUT | WHO I |
| **CH-157MN** | 273 | >5000 | 2582 | >5000 | NF2 MUT | WHO III |
| **GRA** | 66 | 495 | 4989 | >5000 | NF2 MUT | WHO I |
| **KT21-MG1** | 108 | 139 | 713 | 4236 | NF2 MUT | WHO III |
| **Patient 9** | 661 | 3475 | >5000 | >5000 | NF2 MUT | WHO I |
| **Patient 13** | 111 | 434 | 181 | 665 | NF2 MUT | WHO I |
| **Patient 16** | 170 | 3981 | >5000 | >5000 | NF2 MUT | WHO I |
| **Patient 17** | 210 | 1148 | 4498 | >5000 | NF2 MUT | WHO II |
| **Patient 18** | 74 | 332 | 4592 | >5000 | NF2 MUT | WHO I |
| **Patient 19** | 261 | >5000 | >5000 | >5000 | NF2 MUT | WHO II |
| **Patient 20** | 180 | 1941 | 3556 | >5000 | NF2 MUT | WHO I |
| **Pro** | 220 | 457 | 2113 | 3459 | NF2 MUT | n/a |
| **ARC** | 51 | 257 | 1005 | 2148 | NF2 WT | WHO I |
| **FAS** | 77 | 352 | 334 | 444 | NF2 WT | WHO I |
| **GAB** | 286 | >5000 | 3899 | >5000 | NF2 WT | WHO I |
| **IOMM-Lee** | 351 | >5000 | >5000 | >5000 | NF2 WT | WHO III |
| **MUL** | 56 | 101 | 499 | 1936 | NF2 WT | WHO I |
| **NOR** | 131 | 500 | 453 | 4966 | NF2 WT | WHO I |
| **Patient 5** | 187 | 1479 | 4797 | >5000 | NF2 WT | WHO II |
| **Patient 6** | 1503 | >5000 | >5000 | >5000 | NF2 WT | WHO I |
| **Patient 12** | 1199 | >5000 | >5000 | >5000 | NF2 WT | WHO I |
| **Patient 22** | 58 | 76 | 1050 | 667 | NF2 WT | WHO II |
| **Patient 8** | 673 | >5000 | >5000 | >5000 | n/a | WHO I |
| **Patient 23** | 166 | 710 | n.d. | >5000 | n/a | n/a |
| **Patient 26** | 1059 | 4227 | n.d. | >5000 | n/a | n/a |

We treated three resistant and six sensitive cell lines with the four TEAD inhibitors (IAG-933 and MYF-03-176 (500 nM), GNE-7883 (5000 nM), K-975 (1000 nM), all within the on-target ranges of all inhibitors ^38–41^) for 72 hours and isolated total RNA (n = 3 per cell line and inhibitor). We detected a significant downregulation of canonical YAP1/TAZ targets (CTGF, CYR61, ANKRD1, AMOTL2) upon TEADi treatment via qRT-PCR in all TEADi-treated cell lines (Figure 2D; Suppl. Figure 2G), regardless of the NF2 status or sensitivity/resistance to TEADi. In TEAD-sensitive cell lines, the ability of a specific inhibitor correlated with the degree of the downregulation of YAP1/TAZ-TEAD targets, e.g. IAG-933 and to a lesser degree MYF-03-176 resulted in the most significant decrease of target gene expression. Surprisingly, TEAD-resistant cell lines frequently displayed a larger degree of downregulation of YAP1/TAZ-TEAD target genes, suggesting that resistance to TEADi is not caused by the inability of TEADi to suppress specific YAP1/TAZ-TEAD targets, but rather the presence of additional mitogenic pathways.

These results suggest that multiple subtypes of meningioma are reliant on YAP1/TAZ-TEAD signaling. This was further exemplified by the fact that YAP1/TAZ double knockout led to a growth arrest even in TEADi-resistant lines. However, while TEADi-sensitive lines are already responsive to lower TEADi doses, TEADi-resistant cells are not, potentially caused by the presence of other mitogenic pathways that can compensate to a degree.

### TEADi treatment results in a downregulation of YAP1/TAZ-TEAD signaling in both TEADi-sensitive and -resistant cell lines

To analyze the response of meningioma cells to TEADi in more detail, we performed RNA-Seq analysis for five meningioma cell lines (four NF2mut and one NF2 wild type lines) treated with either DMSO or two different TEAD inhibitors (IAG-933 and MYF-03-176, 500 nM each) for 72 hours (n = 3 per condition and cell line). We included three TEADi-sensitive (BenMen1 and BrMen (both low-grade NF2mut), KT21-MG1 (high-grade NF2mut)) and two TEADi-resistant lines (CH-157MN (high-grade NF2mut), IOMM-Lee (NF2wt). In line with our previous results ^30^, we identified a significantly higher baseline expression of YAP1/TAZ-TEAD target genes (CCN1, CCN2, ANKRD1, AMOTL2) in low-grade NF2mut cell lines, compared to high-grade NF2mut or NF2wt cell lines (Suppl. Figure S3A). Differential gene expression analysis followed by Principal Component Analysis (PCA) and hierarchical clustering of the DMSO-treated conditions revealed a clustering broadly along the lines of TEADi-sensitive cell lines (BenMen1, BrMen, KT21-MG1) and TEADi-resistant lines (CH-157MN, IOMM-LEE) (Figure 3A). TEADi-treated conditions remained in these clusters, suggesting that the overall cell line state is dominant over the transcriptional changes induced by TEADi treatment (Figure 3B). However, TEADi-treated cells displayed a significant shift away from DMSO-treated cells that was consistent across all cell lines. This degree of this shift correlated with the degree of inhibition of YAP1/TAZ-TEAD signaling and was greater for IAG-933-treated versus MYF-03-176-treated cells (Figure 3B-D, Suppl. Figure S3B).

**Figure 3:**
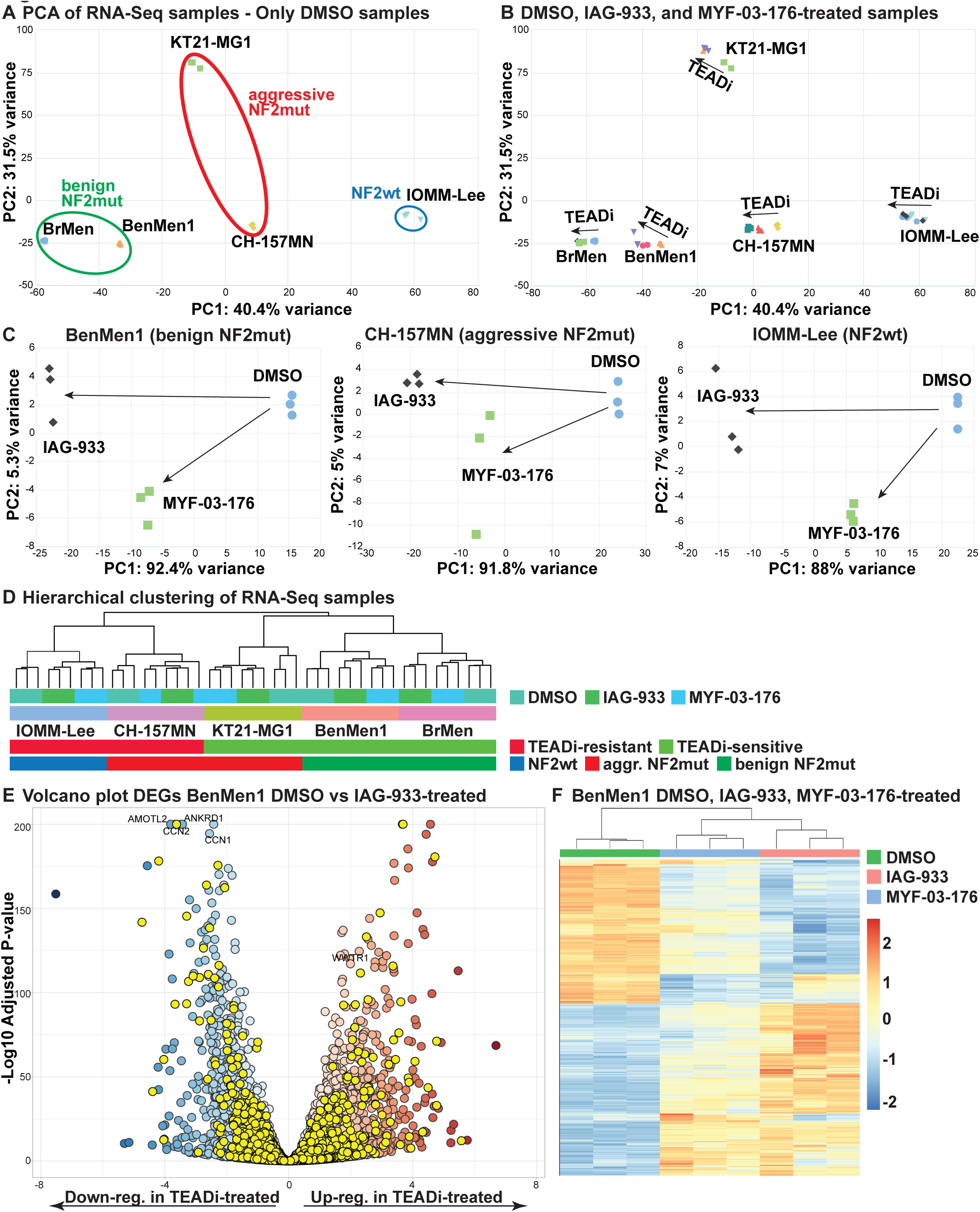
TEADi treatment results in a downregulation of YAP1/TAZ-TEAD signaling in both TEADi-sensitive and -resistant cell lines. A) PCA plot with RNA-Seq data of DMSO-treated samples of benign NF2mut (BenMen1, BrMen), aggressive high-grade NF2mut (KT21-MG1, CH-157MN), and NF2wt (IOMM-Lee) meningioma cell lines (n = 3 each). B) PCA plot showing DMSO, IAG-933, or MYF-03-176-treated samples of the same six meningioma lines (n = 3 each). C) PCA plots showing DMSO, IAG-933, or MYF-03-176-treated samples of benign NF2mut (BenMen1, left), high-grade NF2mut (CH-157MN, middle), or NF2wt (IOMM-Lee, right panel) meningioma cell lines. D) Hierarchical clustering of all five cell lines shows clustering along the lines of NF2 status and TEADi-sensitivity/resistance. E) Volcano plot showing differentially expressed genes between BenMen1 cells treated with DMSO vs TEADi (IAG-933). Yellow dots represent known YAP1/TAZ-TEAD target genes. F) Heatmap showing expression of significantly regulated genes in DMSO, IAG-933, and MYF-03-176-treated BenMen1 meningioma cells.

Treatment with both IAG-933 and MYF-03-176 resulted in highly significant changes of gene expression in all five cell lines (adjusted p-value (padj) < 0.05; fold change (FC) > 1.5) (Suppl. Figure S3C). The gene expression changes induced by TEADi (compared to DMSO) significantly overlapped between IAG-933 and MYF-03-176-treated samples, although IAG-933 resulted in a higher number of deregulated genes and a significantly stronger degree of deregulation (Figure 3E-F, Suppl. Figure S3C-F). In addition, transcriptomic changes induced by the TEADi treatment showed highly significant overlaps between the different cell lines (including both resistant and sensitive lines), suggesting that TEADi treatment induces similar transcriptomic changes in both sensitive and resistant lines (Suppl. Table S1). Lastly, the differentially expressed genes were significantly enriched for YAP1/TAZ-TEAD target genes, suggesting that a significant portion of the TEADi-induced transcriptomic changes are mediated by on-target inhibition of YAP1/TAZ-TEAD signaling (Suppl. Table S1). In summary, TEADi treatment induces highly significant changes in the transcriptome of meningioma cells. These changes were highly similar between sensitive and resistant lines, and the degree of gene regulation correlates with inhibitor efficacy.

### TEADi induces highly similar transcriptional changes compared to YAP1/TAZ knockout

To compare the transcriptional changes induced by TEADi treatment (IAG-933, MYF-03-176) versus combined knockout of YAP1 and TAZ, we performed RNA-Seq from CRISPR-Cas9-edited (CD8A; combined knockout of YAP1 and TAZ (Y+T KO)) BenMen1 and KT21-MG1 cells (Suppl. Table S2) and compared the transcriptional programs to those observed upon TEADi treatment. Total RNA was isolated 72 hours post-gene editing. PCA plots revealed similar gene expression patterns between Y+T KO cells and TEADi-treated cells for both BenMen1 and KT21-MG1 cells (Figure 4A, Suppl. Figure S4A). While the two control conditions (CD8A KO, DMSO-treated) clustered closely together, IAG-933- and MYF-03-176-treated samples clustered closer to Y+T KO samples. Of note, for BenMen1 cells, Y+T KO cells clustered together with IAG-933 samples, whereas for KT21-MG1 cells, Y+T KO samples clustered separately from IAG-933 samples, although adjacent. MYF-03-176-treated samples clustered between control and Y+T KO and IAG-933-treated samples. Heatmaps showing the expression of 2212 YAP1/TAZ target genes in control, TEADi, and Y+T KO samples, displayed a highly similar regulation of YAP1/TAZ-TEAD target genes between IAG-933-treated and Y+T KO samples, while MYF-03-176-treated samples again showed a regulation between control and Y+T KO samples (Figure 4B). We analyzed the expression of the known YAP1/TAZ-TEAD target genes *CCN1*, *CCN2*, *ANKRD1*, and *AMOTL2* (Figure 4C, Suppl. Figure S4B-C), and found that in BenMen1 cells, IAG-933 treatment and combined Y+T KO resulted in the same degree of downregulation of these targets. By contrast, in KT21-MG1 cells, combined Y+T KO resulted in a significantly stronger downregulation of these same target genes compared to IAG-933 treatment. In both cell lines, MYF-03-176 treatment resulted in a less significant inhibition of YAP1/TAZ-TEAD target gene expression. Lastly, we observed highly significant overlaps between the DEGs induced by TEADi-versus DMSO-treated and Y+T KO versus CD8A KO conditions (absolute FC of 1.5; FDR < 0.05) (Figure 4D, Suppl. Figure S4D). In BenMen1 and KT21-MG1 cells we observed that 72% (3556 out of 4940) and 70% (5260 out of 7568) DEGs induced by Y+T KO (compared to CD8A KO) were also differentially regulated in IAG-933-treated samples (compared to DMSO control). Using a Fisher’s exact test to analyze the overlap between the top 1000 DEGs, we observed a highly statistically significant overlap between the transcriptional changes induced by YAP1+TAZ KO and IAG-933/MYF-03-176 treatment responses in both BenMen1 and KT21-MG1 cell lines (FDR < 1e-300). Taken together, these data suggest a shared transcriptional program between genetic YAP/TAZ loss and pharmacological TEAD inhibition. However, the degree of transcriptional regulation can depend on both the cell line and the inhibitor (e.g. treatment with IAG-933 induced more dramatic changes).

**Figure 4:**
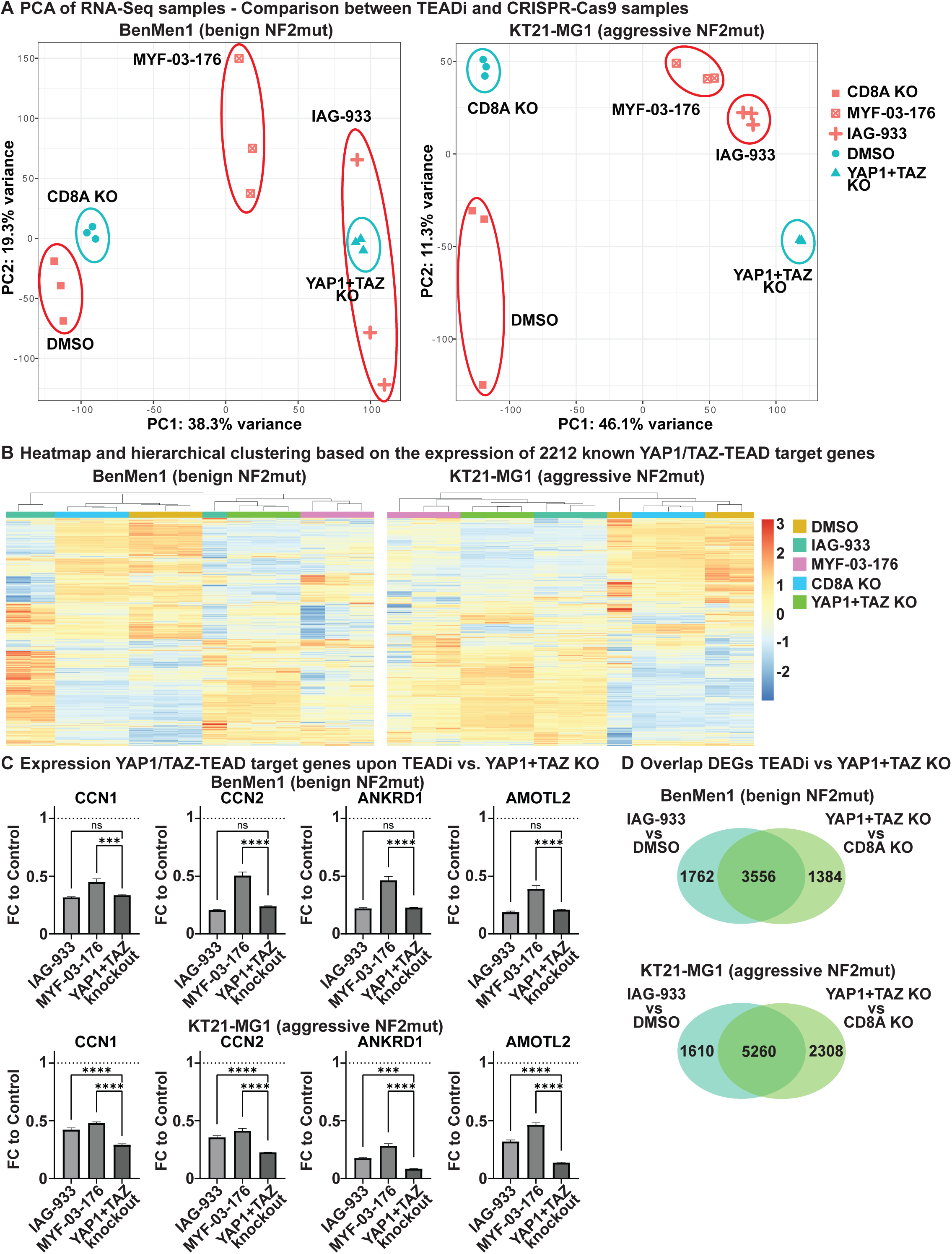
TEADi induces highly similar transcriptional changes compared to YAP1/TAZ knockout. A) PCA plots comparing DMSO-treated, TEADi-treated (IAG-933, MYF-03-176), CD8A KO, and YAP1+TAZ knockout BenMen1 (left) and KT21-MG1 (right) cell lines. B) Heatmaps showing the expression of 2212 YAP1/TAZ target genes in the same sample groups. C) Expression of the YAP1/TAZ target genes CCN1, CCN2, ANKRD1, and AMOTL2 in TEADi-treated and Y+T KO samples from BenMen1 and KT21-MG1 cell lines. Graphs show the fold change (FC) compared to their respective controls (DMSO-treated for IAG-933 and MYF-03-176-treated samples and CD8A KO for Y+T KO). D) Overlap of DEGs from IAG-933-treated samples (compared to DMSO) and Y+T KO samples (compared to CD8A KO) for BenMen1 and KT21-MG1 cell lines. All experiments were performed in triplicates. Error bars show SD (C). Analysis was done using an ordinary one-way ANOVA (C) with multiple comparisons testing.

### TEADi induces MEK-ERK and mTOR-S6 (but not PI3K-AKT) signaling in TEADi-resistant cell lines

KEGG pathway analysis of RNA-Seq data derived from TEADi-versus DMSO-treated cells revealed a significant enrichment of terms related to “DNA replication” and “Cell Cycle” (Suppl. Table S3). In addition, we observed a significant enrichment of terms related to RTK-PI3K, RTK-Ras-PI3K, RTK-Ras-ERK, FAK, RhoA, BMP, JAK-Stat, and Wnt signaling pathways for multiple cell lines. To confirm the differential regulation of the PI3K and MAPK pathways upon TEADi-treatment, we treated seven TEAD-sensitive (BenMen1, BrMen, KT21-MG1, ARC, FAS, MUL, patient tumor 16) and two TEADi-resistant lines (CH-157MN, IOMM-Lee) with IAG-933 and MYF-03-176 (500 nM each) for 72 hours, isolated total protein, and performed Western Blots for phospho- and total levels of AKT, S6, and ERK (Figure 5A, Suppl. Figure S5A). We observed a significant decrease in phospho-S6 levels in all TEADi-sensitive lines upon treatment with both IAG-933 and MYF-03-176, with the former showing a greater degree of regulation. In addition, we also observed a significant decrease in phospho-ERK levels in most TEADi-sensitive cell lines upon treatment. By contrast, we observed a significant increase in phospho-S6 levels in all TEADi-resistant lines upon treatment. The degree of increase was generally higher in IAG-933 treated cells, compared to MYF-03-176. Similarly, phospho-ERK levels were also increased in most TEADi-resistant lines upon treatment. In summary, we identified diametrically opposed activation of MAPK and mTOR-S6 pathways in TEADi-sensitive versus -resistant meningioma lines upon TEADi treatment.

**Figure 5:**
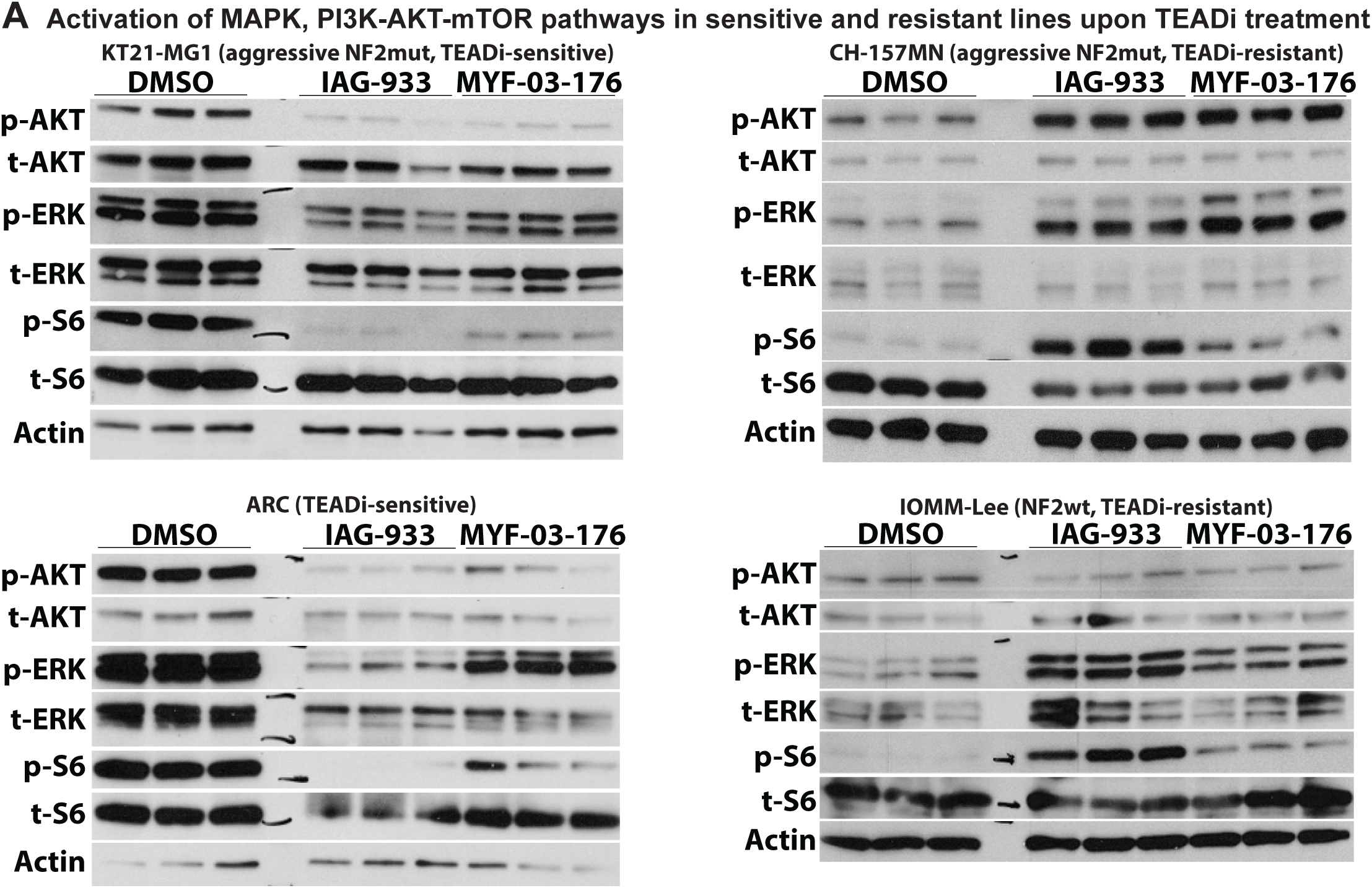
TEADi induces MEK-ERK and mTOR-S6 (but not PI3K-AKT) signaling in TEADi-resistant cell lines. A) Representative western blots showing phosphorylated and total protein levels of AKT, ERK, and S6 in two TEADi-sensitive (KT21-MG1, ARC) and two TEADi-resistant (CH-157MN, IOMM-Lee) meningioma cell lines upon treatment with either DMSO (control) or TEADi (IAG-933 or MYF-03-176, 500 nM each). Actin was used as a loading control.

### Co-targeting MAPK and mTOR-S6 signaling is able to overcome resistance to TEADi

Since we observed a significant upregulation of MEK-ERK and mTOR-S6 pathways upon TEADi treatment of TEADi-resistant meningioma cell lines, we next assessed whether cotreatment of resistant cells with MEK (MEKi (trametinib)), PI3K (PI3Ki (pictilisib)), or mTOR inhibitors (mTORi (torin-2)) can increase the efficacy of TEADi in these cells. We treated TEADi-sensitive and -resistant cell lines with either IAG-933 (200 nM (sensitive lines), 500 nM (resistant lines)), MEKi (1, 5, 10 nM), or mTORi (5, 10, 20 nM), alone or in combination. Treatment efficacy was measured via live cell imaging and endpoint Cell Titer-Glo measurements. While co-treatment with MEKi or mTORi did not significantly increase the efficacy of TEADi in TEADi-sensitive cell lines, it was able to significantly increase the efficacy of TEADi in all tested resistant cell lines, even at low MEKi (1 nM) or mTORi (5 nM) concentrations (Figure 6A-B, Suppl. Figure S6A-B). Importantly, the combination of TEADi and MEKi or mTORi showed a significantly greater inhibition of growth and cell viability than MEKi or mTORi alone, or the combination of MEKi and mTORi (without TEADi) in all tested TEADi-resistant cell lines (Figure 6A-B, Suppl. Figure S6A-B). Treatment with a PI3K inhibitor (Pictilisib, 50-100 nM) did not show any significant efficacy (Figure 6A).

**Figure 6:**
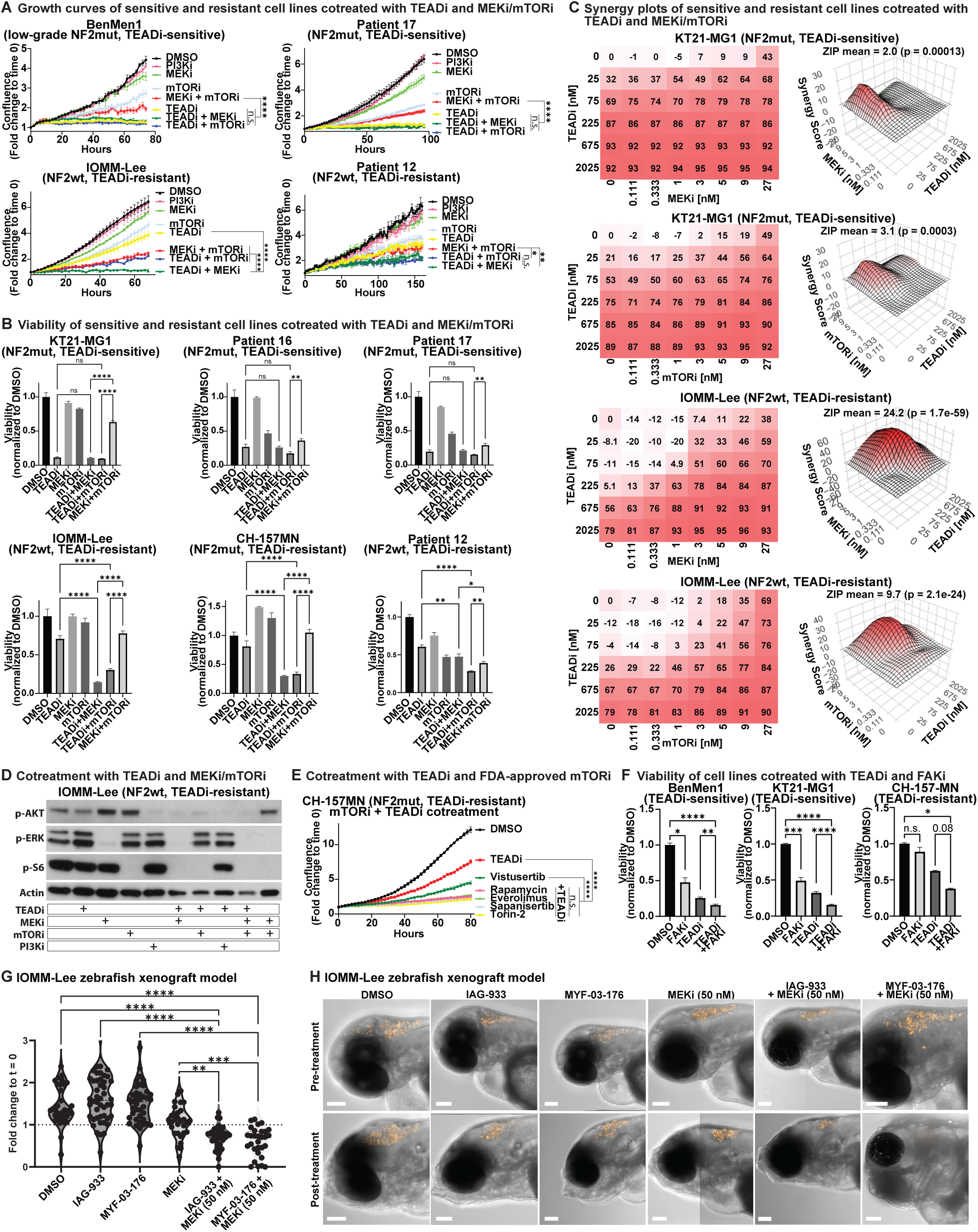
Co-targeting MAPK, mTOR-S6, or FAK signaling is able to overcome resistance to TEADi. A-B) Growth curves (A) and end point viability (B) of TEADi-sensitive or -resistant meningioma cell lines upon co-treatment with combinations of TEADi (IAG-933, 200 nM (sensitive lines) or 500 nM (resistant lines)), PI3Ki (pictilisib, 50 nM), MEKi (trametinib), or mTORi (torin-2). All experiments were performed in triplicates. C) Synergy plots for TEADi/MEKi or TEADi/mTORi combination treatments of TEADi-sensitive (KT21-MG1) or -resistant (IOMM-Lee) meningioma cell lines. D) Western Blot showing activation of MAPK (p-ERK), PI3K-AKT (p-AKT), or mTOR-S6 (p-S6) pathways in TEADi-resistant IOMM-Lee cells upon treatment with TEADi (IAG-933), MEKi (trametinib), mTORi (torin-2), or PI3Ki (pictilisib) in various combinations. Actin was used as a loading control. E) Growth curves of TEADi-resistant CH-157MN cells upon treatment with TEADi (IAG-933, 200 nM) or additional mTOR inhibitors (torin-2, rapamycin, vistusertib, everolimus, sapanisertib, 10 nM each). All experiments were performed in duplicates. F) End point viability of TEADi-sensitive and -resistant cell lines upon treatment with TEADi (IAG-933, 200 nM) or FAKi (TAE226, 500 nM). G) Violin plot showing fold changes of IOMM-Lee tumors grown in zebrafish larvae upon treatment with DMSO, IAG-933, MYF-03-176, trametinib, or combinations thereof. H) Representative images of RFP+ IOMM-Lee tumor-bearing zebrafish larva before and after 48 hours treatment with indicated compounds (IAG-933 (25 µM), MYF-03-176 (5 µM), MEKi (trametinib, 50 nM)). Scale bars indicate 100 µm. Error bars show SEM (A, B, E, F). Analysis was done using an ordinary one-way ANOVA (B,F,G) or ordinary two-way ANOVA (A, E, comparison at last time point) with multiple comparisons testing. Synergy analysis for two-way treatments was done with SynergyFinder3.0.

We performed additional in vitro treatment experiments followed by CellTiter-Glo viability readout after 5 days of treatment, and used SynergyFinder3.0 to identify synergistic drug effects between TEADi and MEKi or mTORi in TEADi-resistant and -sensitive cell lines. We treated five TEADi-sensitive (ARC, BenMen1, KT21-MG1, patient 5, patient 16), and four TEADi-resistant (IOMM-Lee, CH-157MN, GAB, patient 12) cell lines with increasing concentrations of IAG-933 (25, 75, 225, 675, 2025 nM) and trametinib or torin-2 (0.11, 0.33, 1, 3, 5, 9, 27 nM for both). For TEADi-sensitive cell lines, we only observed synergistic effects between IAG-933 and trametinib at low IAG-933 concentrations (25 and 75 nM), but not at higher concentrations (Figure 6C, Suppl. Figure S6C). By contrast, in TEADi-resistant cell lines we observed synergistic effects with as low as 1 nM of trametinib treatment (Figure 6C, Suppl. Figure S6C). Similarly, cotreatment with torin-2 and IAG-933 in TEADi-sensitive cells (ARC, KT21-MG1, patient 5, patient 16) also only resulted in synergistic effects at low IAG-933 concentrations (25 and 75 nM), whereas cotreatment of TEADi-resistant cells (IOMM-Lee, CH-157MN, GAB, patient 12) revealed synergistic effects for torin-2 concentrations of 1 nM and higher (Figure 6C, Suppl. Figure S6D).

We treated TEADi-resistant IOMM-Lee (NF2wt) and CH-157MN (high-grade NF2mut) cells with IAG-933 (500 nM), trametinib (10 nM), torin-2 (10 nM), or pictilisib (50 nM) and combinations thereof for 72 hours and subsequently isolated whole protein lysates. We performed Western Blots for phospho-AKT, phopsho-S6, and phospho-ERK to assess on-target drug efficacies and observed inhibitor-specific inhibition of these pathways. In IOMM-Lee cells, cotreatment with IAG-933 and trametinib resulted both in a reduction of phospho-ERK and phsopho-S6 levels, demonstrating the direct link between MEK-ERK signaling and PI3K-AKT-independent mTOR activation in these cells upon TEADi treatment (Figure 6D). Similar observations were made in CH-157MN cells (Suppl. Figure S6E).

In summary, combination treatments of IAG-933 and MEK or mTOR inhibitors show significant efficacy in overcoming the short-term resistance to TEADi in TEADi-resistant meningioma cell lines. Importantly, the combination of TEADi with MEK or mTOR inhibitors was superior to either inhibitor alone and was also superior to the combination of MEKi and mTORi.

### Additional FDA-approved mTOR inhibitors show a similar beneficial effect in combination with TEADi

While trametinib has been approved by the FDA for the clinical application in multiple clinical settings, including for use in meningioma and other CNS tumors (such as low-grade glioma in combination with the Raf inhibitor dabrafenib), torin-2 is not an FDA-approved inhibitor and is not currently being assessed in clinical trials, despite high efficacy in multiple preclinical studies ^43^. To this end, we assessed the ability of additional FDA-approved mTOR inhibitors (vistusertib, everolimus, rapamycin, and sapanisertib, with torin-2 as a positive control) to significantly enhance the efficacy of TEADi in TEADi-resistant meningioma cell lines. We treated TEADi-resistant CH-157MN, KT21-MG1,and IOMM-Lee meningioma cells with IAG-933 (200 nM (CH-157MN, KT21-MG1), 500 nM (IOMM-Lee), mTOR inhibitors (1, 3, 5, 10, 20, 50 nM), or a combination of IAG-933 and mTORi and monitored cell growth via live-cell imaging and generated dose-response curves (Figure 6E, Suppl. Figure S6F). As previously observed for torin-2, single-agent treatment with the different mTOR inhibitors alone did not result in a significant growth arrest or reduction in viability (Suppl. Figure S6F-G), apart from 20 and 50 nM doses of torin-2 and sapanisertib. By contrast, co-treatment with IAG-933 and different mTOR inhibitors showed a significant growth reduction beyond the effect of single-agent IAG-933 or single-agent mTORi. In combination with TEADi, Torin-2, sapanisertib, rapamycin, and everolimus displayed the greatest ability to inhibit the growth and viability of these cell lines, whereas vistusertib showed the lowest efficacy. In summary, cotreatment with the FDA-approved mTOR inhibitors everolimus, rapamycin, and sapanisertib is able to significantly increase the efficacy of TEADi against TEADi-resistant meningioma cell lines in a similar manner to torin-2.

### Combined TEAD and FAK inhibition exert additive inhibitory effects in a subset of meningioma cell lines

Since the KEGG terms of multiple cell lines analyzed by RNA-Seq also showed an enrichment of Focal Adhesion signaling in TEADi-treated cell conditions, we assessed the efficacy of a FAK inhibitor (TAE226, 500 nM) alone or in combination with IAG-933 (100 nM for BenMen1 and KT21-MG1; 500 nM for the remaining cell lines) in eight cell lines, including TEADi-sensitive (BenMen1, KT21-MG1) and -resistant cell lines (CH-157MN, IOMM-Lee, Patient 6, 8, 9, 12), and assessed the impact on cell viability via CellTiter-Glo. Patient cell lines 8 and 9 were only tested for single-agent TAE226. Single-agent TAE226 treatment was able to significantly inhibit the viability and growth of five out of nine cell lines (BenMen1 and KT21-MG1 (both TEADi-sensitive NF2mut cell lines); patient tumors 6, 9, and 12), but was generally less effective compared to IAG-933 treatment (Figure 6F, Suppl. Figure S6H). Cotreatment of IAG-933 and TAE226 was able to further significantly reduce the viability compared to the single-agent IAG-933 conditions in two out of seven cell lines (BenMen1, KT21-MG1). CH-157MN and patient tumor 6 cells showed a non-significant trend towards reduced viability in TAE226+IAG-933 co-treated cells, compared to single-agent IAG-933-treated cells (p = 0.08 and 0.07, respectively). In summary, cotreatment of TEADi with FAK inhibitors is beneficial for a subset of meningioma cell lines, including both TEADi-sensitive and -resistant lines. Although the combination with FAK inhibition is less potent compared to MEKi and mTORi cotreatment in TEADi-resistant lines.

### Cotreatment with TEADi and MEKi significantly reduces meningioma tumor growth in a zebrafish xenograft meningioma model

To study the effect of TEADi single and combination treatment in an in vivo model, we developed a zebrafish meningioma xenograft model by transplanting RFP-expressing IOMM-Lee cells into the hindbrain ventricle of 2 days post-fertilization transparent zebrafish larvae (*mitfa^b^*^692^*^/b^*^692^). Tumor volume was quantified via RFP fluorescence 24 hours after tumor cell injection, and zebrafish larvae were subsequently incubated with either DMSO (control; n = 34), IAG-933 (25 µM; n = 46), MYF-03-176 (5 µM; n = 39), trametinib (50 nM; n = 30) (all at pre-determined no observed adverse effect concentrations (NOEC), except for IAG-933 (NOEC > 100 µM)), or combinations thereof (IAG-933 (25 µM) + trametinib (50 nM) (n = 36); MYF-03-176 (5 µM) + trametinib (50 nM) (n = 30). Treatment experiments were performed as three independent sets of experiments (each including similar numbers of all treatment groups). Post-treatment tumor volume was measured 48 hours after treatment start and tumor volume change was calculated compared to the pretreatment volume.

In line with our in vitro results, we observed that IAG-933, and MYF-03-176 alone had no significant effect on in vivo tumor growth compared to DMSO. While DMSO treated tumors grew on average by 44.6% (±7.8% SEM) over the 48 hours’ experimental time frame, IAG-933 (+61.5% ±8.1 %) and MYF-03-176 (+51% ±7.3%) grew to a similar degree (p > 0.5). Trametinib-treated tumors (+7% ±6.9%) grew significantly slower (p = 0.007 to DMSO-treated), but regression was not observed. By contrast, tumors treated with combinations of TEADi (IAG-933 or MYF-03-176) plus trametinib regressed within the 48 hours of treatment (IAG-933 plus 50 nM trametinib (−29.2% ±4.4%) and MYF-03-176 plus 50 nM trametinib (−40.2% ±6.2%), both p < 0.0001 to DMSO-treated) (Figure 6G-H).

In summary, the combination of TEADi with MEKi was able to induce significant decreases in tumor volume within 48 hours in an in vivo zebrafish meningioma xenograft model.

### Whole exome sequencing identifies Ras-Raf pathway mutations in TEADi-resistant cell lines

To identify genetic variants (somatic variants (SV), copy number variation (CNV) analysis) in TEADi-sensitive and resistant meningioma cell lines, we performed whole exome sequencing (WES) of 19 cell lines (Suppl. Table S4A). WES confirmed NF2 status in all tested cell lines apart from PRO, GRA, and patient tumor 19 lines. We observed the highest number of CNVs in KT21-MG1 (sensitive, 848 altered genes), CH-157MN (resistant, 627 altered genes), patient tumor 19 (resistant, prior radiation, 187 altered genes), GAB (resistant, 170 altered genes), and IOMM-Lee (resistant, 143 altered genes), followed by patient tumor 9 (resistant, 84 altered genes), BenMen1 (sensitive, 81 altered genes), and patient tumor 20 (sensitive, 76 altered genes). While TEADi resistant tumors were enriched for a higher degree of genome instability, most sensitive cell lines (patient 18 (21 altered genes), patient 16 (15 altered genes), patient 5 (5 altered genes), NOR (3 altered genes), BrMen (1 altered gene), MUL (1 altered gene), ARC, FAS, GRA, patient 22, PRO (all zero altered genes)) harbored few or no gene amplifications/deletions (Figure 7A; Suppl. Table S4A). Frequent CNVs in addition to chr22 loss included gains of chr3q, chr5, chr8q (four cell lines each), as well as loss of chr1p and chr4 (four cell lines each). Deep CDKN2A/B deletions were observed in three cell lines (CH-157MN, IOMM-Lee, GAB (all resistant)). Specific copy number variations in MAPK pathway components were observed in both TEADi-resistant and sensitive cell lines, including amplifications in Ras and Raf genes, as well as deletions in NF1 and Ras genes (Suppl. Table S4A). A PTEN deletion was observed in KT21-MG1 cells, although these cells harbor residual PTEN expression ^35^.

**Figure 7:**
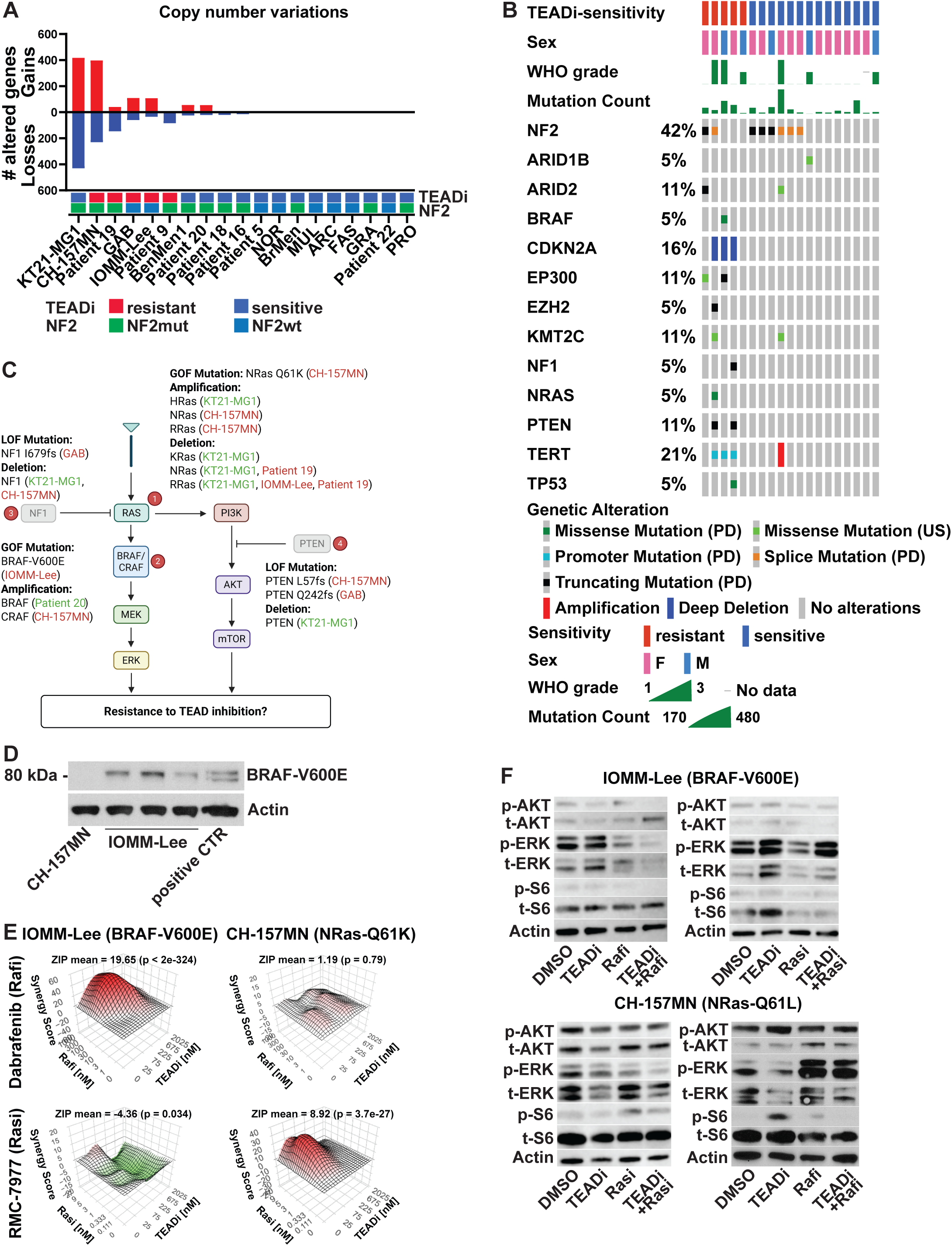
Whole exome sequencing identifies Ras-Raf pathway mutations in TEADi resistant cell lines. A) CNV altered genes, TEADi-response, and NF2 status for all 19 cell lines that underwent WES. B) Oncoprint plot showing somatic variants identified in meningioma cell lines that underwent WES. PD = putative driver, US = unknown significance C) Schematic showing SV and CNV alterations within the MAPK and PI3K-mTOR pathways identified in meningioma cell lines that underwent WES. D) Western Blot for BRAF-V600E of CH-157MN, IOMM-Lee, and human BRAF-V600E-expressing mouse glioma cells (positive control). E) Synergy plots for TEADi/Rafi or TEADi/Rasi combination treatments of BRAF-V600E-(IOMM-Lee) or NRas-Q61K-positive (CH-157MN) meningioma cell lines. F) Western Blots with IOMM-Lee and CH-157MN cell lysates for phospho-and total versions of AKT, ERK, and S6 upon treatment with TEADi, Rasi, Rafi, or combinations thereof. Synergy analysis for two-way treatments was done with SynergyFinder3.0.

Somatic variants/point mutations were observed in all cell lines (Figure 7B-C; Suppl. Table S4B). We observed meningioma-typical mutations in the two NF2wt patient tumor cell lines (patient tumor 5: BAP1 (splice region variant); patient tumor 22: SMO1 (L412F), both TEADi-sensitive). We observed multiple mutations potentially affecting MAPK and PI3K-mTOR pathway activation in TEADi-resistant cells, including GOF mutations in BRAF (V600E) in IOMM-Lee cells and NRas (Q61K) in CH-157MN cells, as well as LOF mutations in NF1 (I679framshift (fs)) in GAB cells, and PTEN in CH-157MN (L57fs) and GAB (Q242fs) cells. We also identified an NF1 splice region variant/intron variant in KT21-MG1 cells, it is unknown whether this type of mutation results in a functional impairment/loss of NF1 in these cells. KT21-MG1 cells also experienced loss of KRas, NRas, and RRas, but gain of HRas, and it is unknown how this would affect the impact of an NF1 loss-of-function mutation. We furthermore observed multiple potential LOF mutations in chromatin-remodeling genes, including ARID1B (intron variant; patient tumor 9), ARID2 (Q1194stop; patient tumor 9), EZH2 (Q194stop; CH-157MN), and EP300 (C1621stop; IOMM-Lee). KMT2C loss-of-function (LOF) mutations were observed in three NF2wt cell lines (IOMM-Lee, GAB, NOR) and were previously reported to suppress NF2 expression ^44^. Potential RB1 LOF mutations (all intronic variants) were detected in 13 cell lines (including sensitive and resistant lines), TERT gain-of-function (GOF) promoter mutations (−124 bp (C228T)) were detected in three TEADi-resistant cell lines (CH-157MN, IOMM-Lee, GAB), and a TP53 LOF R273H was detected in one TEADi resistant cell line (GAB).

We validated the presence of the BRAF-V600E mutation in IOMM-Lee cells via Western Blot (Figure 7D). We then performed in vitro co-treatment experiments with IOMM-Lee (BRAF-V600E) and CH-157MN (harboring a NRas-Q61K mutation) cells, treating these cells with TEADi (IAG-933, same concentration range as above) and either Dabrafenib (BRAF-V600E inhibitor) (1, 3, 10, 30, 100, 300, or 1000 nM) or RMC-7977 (pan-Ras inhibitor) (0.111, 0.333, 1, 3, 5, 9, or 27 nM), followed by CellTiter-Glo viability readout, and used SynergyFinder3.0 to identify synergistic drug effects. We observed that cotreatment of IOMM-Lee cells with TEADi and Dabrafenib (p < 2 x 10^-324^) but not with TEADi and RMC-7977 (p = 0.034) showed highly significant synergistic effects. Conversely, co-treatment of CH-157MN cells with TEADi and RMC-7977 (p = 3.7 x 10^-27^) but not with Dabrafenib (p = 0.79) showed significant synergistic effects (Figure 7E). Western Blot analysis of phospho- and total levels of AKT, S6, and ERK confirmed the treatment-specific inhibition of these pathways (Figure 7F).

These results suggest that TEADi resistant cell lines are enriched for a higher degree of genome instability and frequently harbor gain- or loss-of-function mutations in the Ras-Raf pathway. WES also confirms that NF2wt cell lines are by default not less sensitive to TEADi compared to NF2mut cells.

## Discussion

Although the majority of meningiomas are benign and curable by surgery alone, a significant proportion are of higher grade, display aggressive growth behavior, and invade into the brain and recur even after multiple surgeries. In addition, a significant portion of meningiomas grows in locations that are difficult to access via surgery due to their deep location and/or their frequent involvement of major blood vessels and cranial nerves (such as the skull base and optic nerves), preventing complete resection and leading to recurrences and/or worsening symptoms even in low-grade tumors. Chemotherapeutic drugs commonly used to treat other solid tumors—such as cyclophosphamide, doxorubicin, temozolomide, and vincristine—have generally shown limited effectiveness in meningiomas ^8–12^. Meanwhile, several targeted therapies, including inhibitors of FAK, PI3K, CDK4/6, and EZH2, are currently being evaluated in clinical trials and may demonstrate benefit in a subset of patients ^17^.

The clear biological convergence of multiple molecular meningioma subtypes on YAP1/TAZ-TEAD-dependent signaling serves as a strong argument for the use of TEAD inhibitors in meningioma. NF2 loss, the most common genetic event in meningioma, disrupts Hippo pathway regulation and promotes deregulated YAP1/TAZ activity, while recurrent YAP1 fusions in NF2wt tumors produce constitutively active TEAD-driven transcriptional programs. This convergence suggests that TEAD represents a shared downstream vulnerability across genetically distinct but mechanistically related meningioma subgroups and provides a strong rationale for inhibiting the YAP1/TAZ-TEAD complex in meningioma. 1) Our experimental data suggest that both NF2mut and wild type meningioma cells rely on YAP1/TAZ-TEAD signaling for their growth, since we observed a highly significant reduction in cell growth upon combined CRISPR-Cas9-mediated genetic perturbation of YAP1 and TAZ. 2) We observed that treatment with TEAD inhibitors by itself was sufficient to significantly reduce the growth of multiple “TEADi-sensitive” meningioma lines (predominantly lower-grade NF2mut and NF2wt cell lines as well as a subset of high-grade NF2mut lines). 3) We observed that NF2wt meningioma cells are as sensitive to TEADi as NF2mut lines in the absence of additional resistance-conferring mutations. 4) We found that TEADi-resistant cell lines showed a higher degree of genomic rearrangements compared to sensitive cell lines (with the exception of KT21-MG1 cells) and frequently harbored activating mutations in the MAPK pathway. 5) We found that resistant cells respond to TEADi upon targeting compensatory pathways with MEK or mTOR inhibitors. Importantly, the combination of TEADi with MEK or mTOR inhibitors was superior to either inhibitor alone and was also superior to the combination of MEKi and mTORi, suggesting that TEADi sensitizes these cells to MEK/mTOR inhibition. These results are in line with findings from other cancer types ^45^. Synergistic effects between TEADi and MEKi/mTORi were observed at concentrations as low as 1 nM for both the MEK inhibitor trametinib and the mTOR inhibitor torin-2. We could furthermore show that additional, FDA-approved mTOR inhibitors exert a similar ability to significantly increase the efficacy of TEADi in resistant cells. In TEADi-sensitive meningioma cell lines, combination therapies of TEADi and MEKi/mTORi only show synergistic effects at lower IAG-933 doses. The combination of TEADi and MEKi/mTORi in TEADi-sensitive meningiomas may however still be beneficial as it may allow lower doses of TEADi to be effective, which may allow to limit known on-target toxicities of TEAD inhibitors, such as kidney toxicity and proteinuria ^46^. In addition, although the blood brain barrier is generally not an issue for drug delivery into meningiomas, limited in vivo bioavailability of TEAD inhibitors may still reduce drug delivery into the tumor tissue. We validated the in vitro findings in a zebrafish xenograft model and observed significant tumor inhibition upon cotreatment with TEADi (IAG-933, MYF-03-176) and MEKi at non-toxic concentrations, suggesting a therapeutic window for this inhibitor combination.

In addition to MEK and mTOR inhibition, we also identified FAK inhibition as an additional option for cotreatment. Although not as effective as MEKi/mTORi, cotreatment with the FAK inhibitor TAE226 was able to significantly further increase the efficacy of TEADi in around half of the tested cell lines. Combined FAK and MEK inhibition has recently been shown to be effective in a murine model of NF2-related schwannomatosis and FAK inhibition is currently being assessed in clinical trials for meningiomas ^15,47^. RNA-Seq analysis of TEADi-treated meningioma cells identified additional enriched pathways, such as WNT, RhoA, Interleukin2/6-Jak-STAT, and IFN signaling pathways, and future experimentation will be needed to ascertain the importance of these pathways in the resistance to TEADi. Our results suggest that the activation of the mTOR-S6 pathway in TEADi-resistant lines upon TEADi treatment may occur via the ERK pathway, rather than via the PI3K-AKT axis. ERK-dependent mTOR activation can occur via the ERK1/2-mediated phosphorylation of Raptor or TSC2 ^48,49^. However, the factors that contribute to the activation of MAPK signaling in TEADi-resistant cells upon TEADi treatment are still unclear and will need to be explored in future experimentation.

Whole exome sequencing identified several mutations potentially contributing to TEADi resistance, including several mutations within the MAPK pathway. We validated the efficacy of cotreatment with TEADi and Dabrafenib (BRAF-V600E inhibitor) and RMC-7977 (Ras inhibitor) against IOMM-Lee cells (harboring a BRAF-V600E mutation) and CH-157MN cells (harboring an NRas-Q61K mutation). Although BRAF-V600E mutations and BRAF fusions have been described in a small subset of meningiomas ^50,51^, NRas mutations have not been detected in meningiomas. Due to their rarity, it is unlikely that these mutations were present in the parental meningiomas these cell lines were derived from and it is unlikely that this type of mutation would contribute to initial treatment resistance in treatment-naïve meningiomas. However, off-target resistance to TEADi may emerge through these pathways during treatment. In line with our results, using a CRISPR-Cas9 dropout screen, Kulkarni et al have previously identified mutations in the MAPK and PI3K-AKT pathways (including NF1 and PTEN loss) to be involved in resistance to TEADi in mesothelioma ^45^.

We observed significant differences in the efficacy of the four tested TEAD inhibitors (IAG-933, MYF-03-176, GNE-7883, K-975) to inhibit the growth of both TEADi-sensitive and - resistant meningioma cell lines. This was in part due to the higher on-target efficacy of IAG-933 and MYF-03-176, exemplified by their significantly stronger ability to down-regulate YAP1/TAZ-TEAD target genes. However, while all four inhibitors were able to inhibit the growth of TEADi-sensitive cell lines to a degree, only IAG-933 was still able to partially inhibit the growth of TEADi-resistant lines, albeit at a higher concentration. It was puzzling why MYF-03-176 was able to inhibit the growth of TEADi-sensitive lines with a similar efficacy as IAG-933, but performed significantly worse than IAG-933 against resistant lines. Of note, IAG-933 and MYF-03-176 have reported different affinities to the four different TEAD proteins, with MYF-03-176 showing a higher affinity to TEAD1&3 ^40^. The four inhibitors inhibit YAP1/TAZ-TEAD signaling via different mechanisms. MYF-03-176 and K-975 target the TEAD palmitate pocket and irreversibly prevent auto-palmitoylation, GNE-7883 reversibly binds to the lipid pocket of TEADs, whereas IAG-933 binds to and reversibly blocks the TEAD Ω-loop that is necessary for YAP1/TAZ-TEAD interaction. Future experiments will be necessary to determine whether a specific type of TEAD inhibitor is more effective against specific tumor and meningioma subsets.

All of our treatment experiments were limited to short-term treatments (1 week maximum) and the resistance to long-term TEADi treatment in meningiomas will need to be explored. Studies in NF2mut mesothelioma cell lines have identified increased AP-1 (activator protein-1) transcription, MYC and HGF overexpression, as well as upregulation of MAPK (NF1 loss) and JAK-STAT (SOCS3 loss) as potential causes for acquired resistance to TEADi (K-975, GNE-7883, MRK-A) ^39,45,52,53^. A direct combination with MEK inhibitors may be able to overcome or prevent some of these resistance mechanisms from occurring.

Several TEAD inhibitors are currently or have previously been tested in clinical trials. Despite their potential benefit in NF2mut/YAP1-TAZ-driven tumors (such as meningioma), TEAD inhibitors have encountered several hurdles in clinical trials. Both IAG-933 (NCT04857372, Novartis) and VT3989 (NCT04665206, Vivace Therapeutics) have reported limited clinical efficacy in their respective patient population ^42,54^. Co-targeted of TEADi with MEKi/mTORi may yield increased clinical efficacy. Furthermore, TEADi has recently been linked to on-target kidney toxicity ^46^, which could make the current generation of TEADi unsuitable for clinical use. However, the goal of this study is a proof-of-concept to show that TEADi is able to reduce the growth and viability of meningiomas cells. Future research could determine what causes this toxicity and lead to new next-generation inhibitors (that are more selective to a specific TEAD variant) or nanoparticles formulations could be utilized to deliver TEADi more locally.

In summary, our results establish YAP1/TAZ-TEAD signaling as a prominent oncogenic driver in meningioma and serve as a proof-of-concept and strong rational for the clinical application of TAD inhibitors in meningioma. We furthermore explore pathways that contribute to the resistance to TEADi and identify MEK, mTOR, and FAK inhibition as effective combination therapies.

## Methods

### Cell lines

BenMen1 cells were purchased from DSMZ (Deutsche Sammlung von Mikroorganismen und Zellkulturen GmbH). The KT21-MG1, CH-157MN, and IOMM-Lee cell lines were kind gifts of Dr. Jonathan Chernoff (Fox Chase Cancer Center), Dr. Yancey Gillespie (University of Alabama School of Medicine, Birmingham, Alabama), and Dr. Ian McCutcheon (University of Texas, M.D. Anderson Cancer Center, Houston, Texas), respectively. GAB, NOR, FAS, MUL, ARC, PRO, BrMen, and GRA cell lines were previously collected by Dr. Randy Jensen ^55^. Primary patient cell lines were established from patient samples collected under the Huntsman Cancer Institute Total Cancer Care (TCC) IRB (IRB 89989). Primary tumor tissue for the Patient 26 cell line was collected at Emory University under the Core for Brain Tumor, Cerebrospinal fluid, and Plasma Samples IRB protocol (IRB P011587). All tumor tissues from which primary tumor lines were established were graded by a board-certified neuropathologist.

BenMen1 cells were grown in DMEM (Thermo Fisher Scientific 11995065), 20% FBS (Thermo Fisher Scientific A5670701), 1% Pen/Strep). All other cell lines were grown in 10% FBS. For primary cell lines established from primary patient samples, tumor tissue was mechanically dissociated into small pieces, and cultured in DMEM 10% FSB 1% Pen/Strep.

### In vitro growth assays

500-2000 cells per well (depending on the cell line and cell size) were seeded into 96 well plates in their respective media. 24 hours after seeding, 2x inhibitor concentrations were added into the wells to achieve a 1x inhibitor concentration. Appropriate DMSO-only concentrations were used as controls. Each condition was set up in triplicates. Cells were then incubated for 5-7 days (until the DMSO condition reach >80 percent confluency). Live cell imaging was performed using an Incucyte S3 Live-Cell Analysis System with images being acquired every 2 hours. Endpoint viability was measured using the CellTiter-Glo 2.0 Cell Viability Assay (Promega G9241) according to the manufacturer’s instructions. Luminescence was detected using a Microplate Luminometer. For 2-way synergy testing, inhibitors were plated in unicates with concentrations ranging from 25 to 2025 nM (IAG-933) or 0.111 to 27 nM (trametinib, torin-2). Endpoint viability was measured using the CellTiter-Glo 2.0 Cell Viability Assay as described above and 2-way inhibitor synergies were calculated using SynergyFinder 3.0 ^56^.

### Zebrafish xenograft model

All zebrafish experiments were approved under the University of Utah Office of Animal Care and Compliance (OACC) protocol #2585. Homozygous *mitfa* mutation (nacre) zebrafish embryos were grown in E3 embryo medium (5 mM NaCl, 0.17 mM KCl, 0.33 mM CaCl2, 0.33 mM MgSO4) at 28° C until 2 days post-fertilization (dpf). At 2dpf, embryos were dechorionated, anesthetized with 1X tricaine methanesulfonate (MS-222), and aligned in agarose wells. Using a FemtoJet 4i (Eppendorf, Germany) microinjector and glass micropipette, approximately 200 RFP-expressing IOMM-Lee cells were injected into the hindbrain ventricle. Injected embryos were then placed in fresh E3 and incubated at 34° C for 24 hours. Following 24 hours at 34° C, embryos were anesthetized with 1X tricaine and transferred into individual wells of a ZF plate (Hashimoto) for confocal imaging in a CellVoyager CV8000 High-Content Screening System (Yokogawa). Tiled, z-stack images of the entire fish were captured in brightfield (20 ms exposure) and using a 561 nm laser (50% power, 50 ms exposure). Following imaging, all fish were transferred to a 96 well plate with fresh E3. Inhibitors or DMSO control were then added to the E3 of each treatment group at “no observed (adverse) effect concentrations” (NOEC) and the fish were incubated at 34° C for 48 hours. After 48 hours of drug treatment (3 days post-injection, 5 dpf), the fish were anesthetized and transferred into a ZF plate for imaging (see above). Following this second round of imaging, all fish were euthanized. Tumor volume was quantified using CellPathfinder software and fold changes were calculated between pre- and post-treatment images. No observed adverse effect concentrations (NOECs) were determined by titrating compound concentrations and administering to the different groups of non-transplanted nacre larvae at 3 dpf, which were then incubated at 34°C for 48 hours (matching the xenotransplantation timeline). The highest concentration which showed no effect on morphology or behavior was determined to be the NOEC and was used in the xenotransplantation experiments (except for IAG-933, which had a NOEC of > 100 µM).

### Western Blot Analysis

For analysis of downstream pathway activation, cells were seeded into 6 or 12 wells, and incubated with appropriate inhibitor concentrations for 72 hours. Cells were cultured in media containing 0.5% FBS 24 hours prior to and during inhibitor treatment. Cells were incubated with either TEADi (IAG-933, MYF-03-176, 500 nM each), MEKi (trametinib 10 nM), mTORi (torin-2, 10 nM), or PI3Ki (pictilisib, 50 nM). Cells were cultured, lysed, and processed for western blotting by standard methods. Proteins were resolved by SDS/PAGE (NuPAGE 10% Bis/Tris; LifeTech) according to XCell Sure Lock Mini-Cell guidelines, blocked with 5% milk/TBST and probed with specified antibodies overnight at 4°C in 5% BSA/TBST. After three TBST rinses, species-specific secondary antibodies were added in 5% milk/TBST. Blots were rinsed three times with TBST before being developed with Amersham ECL Western Blotting Detection Reagents (GE Healthcare). For a list of antibodies used see Suppl. Table S5A.

### Inhibitors

Inhibitors were purchased from MedChemExpress (MCE), Sigma-Aldrich (SA), or SelleckChem (SC) (IAG-933 (MCE HY-153811; SC E1490); MYF-03-176 (SA SML3686-5MG); GNE-7883 (MCE HY-147214), K-975 (MCE HY-138565); trametinib (MCE HY-10999); torin-2 (MCE HY-13002); pictilisib (MCE HY-50094); vistusertib (MCE HY-15247); everolimus (MCE HY-10218); rapamycin (MCE HY-10219); sapanisertib (MCE HY-13328).

### CRISPR-Cas9 gene editing

Cells were transiently transfected with ribonucleoprotein complexes composed of chemically synthesized 2′-*O*-methyl-3′phosphorothioate-modified single guide RNAs (sgRNAs) and purified Cas9 protein using an Amaxa 4D nucleofector as previously described ^57^. Two guides each were used for YAP1 and WWTR1 (TAZ). A guide RNA targeting CD8A served as a control. Guide sequences are listed in Suppl. Table S5B. Successful gene editing was assessed 96 hours post nucleofection via targeted PCR followed by Sanger Sequencing and subsequent Synthego Inference of CRISPR Edits (ICE) analysis. Primer sequences for targeted PCRs are listed in Suppl. Table S5C Functional impact of YAP1/TAZ codeletion was assessed via qRT-PCR for YAP1/TAZ-TEAD target gene expression and live-cell imaging.

### RNA Isolation, PCR, and RNA Sequencing

For analysis of YAP1/TAZ-TEAD target gene regulation upon TEADi treatment, cell lines were seeded into 6 well plates and treated with TEADi for 72 hours (IAG-933 (500 nM, MYF-03-176 (500 nM), GNE-7883 (5000 nM), K-975 (1000 nM)). For analysis of YAP1/TAZ-TEAD target gene regulation upon CRISPR-Cas9-mediated genetic ablation of YAP1/TAZ, cells were seeded into 6 well plates 48 hours after CRISPR-Cas9 gene editing and RNA was isolated 48 hours later. RNA was extracted using the Qiagen RNeasy Mini Kit according to the manufacturer’s instructions. Genomic DNA was removed by on-column DNase digestion. RNA concentration was measured with a Qubit RNA BR Assay Kit (Fisher Scientific #Q10211). RNA quality was evaluated with an Agilent Technologies RNA ScreenTape Assay (5067-5576 and 5067-5577). Poly(A) RNA was purified from total RNA samples (10-500 ng) using the NEBNext Poly(A) mRNA Magnetic Isolation Module (E7490). Stranded RNA sequencing libraries were prepared as described using the NEBNext Ultra II Directional RNA Library Prep Kit for Illumina (E7760L). Purified libraries were qualified on an Agilent Technologies 4150 TapeStation using a D1000 ScreenTape assay (cat# 5067-5582 and 5067-5583). The molarity of adapter-modified molecules was defined by quantitative PCR using the Kapa Biosystems Kapa Library Quant Kit (cat#KK4824). Individual libraries were normalized to 5 nM in preparation for Illumina sequence analysis. Sequencing libraries were chemically denatured in preparation for sequencing. Following transfer of the denatured samples to an Illumina NovaSeq X instrument, a 151 x 151 cycle paired end sequence run was performed using a NovaSeq X Series 25B Reagent Kit (20104706). For quantitative real-time PCR, RNA was transcribed into cDNA using the SuperScript III kit. PCR experiments were carried out on a Bio-Rad Flex Real-Time PCR System. For primer sequences see Suppl. Table 4C.

### Whole exome sequencing

Genomic DNA from meningioma cell lines was isolated via the PureLink™ Genomic DNA Mini Kit. Library preparation was performed with the NEB NEBNext Ultra II FS DNA Library Prep Kit for Illumina (E7805). Briefly, genomic DNA (100-400 ng) was enzymatically fragmented and NEBNext Unique Dual Index UMI Adaptors (E7395) were ligated to ends of DNA fragments. Size selection was performed with Illumina Purification Beads (20060058) to separate the library into two fractions. A long fraction, ranging from 450 to 800 bp, was retained as a pcr-free library that was sequenced at a depth of 5x whole-genome coverage for copy number analysis. A short fraction, ranging from 250 to 450 bp, was pcr-amplified (5 cycles) and exome enrichment was performed with the Agilent Technologies SureSelect Human All Exon v8 Kit (5191-6891). The exome enriched library was PCR-amplified an additional 8 cycles and libraries were qualified on an Agilent Technologies 4150 TapeStation using a High Sensitivity D1000 ScreenTape assay (cat# 5067-5584 and 5067-5585). The molarity of sequencing libraries was defined by quantitative PCR using the Roche Library Quant Kit Uni (07960140001) and the molarity of individual libraries were normalized to 5 nM in preparation for pooling and sequence analysis. Sequencing libraries were chemically denatured in preparation for sequencing. Following transfer of the denatured samples to an Illumina NovaSeq X instrument, a 151 x 151 cycle paired end sequence run was performed using a NovaSeq X Series 10B Reagent Kit (20085594).

### Bioinformatic Analysis

RNA-seq reads (151 × 151 bp paired-end) were adapter-trimmed with cutadapt and optical duplicates removed with BBMap/Clumpify. Reads were aligned to the GRCh38 reference from Ensembl Release 112 using STAR v2.7.9a and gene-level counts were generated with featureCounts v1.6.3. Downstream analyses were performed in R using DESeq2 (v1.42.1); low-count features (≤5 reads) were filtered prior to testing, and regularized-log (rlog) values were used for visualization. Differential expression was defined at FDR < 0.05 and |log2 fold change| > 0.585 (≈1.5×). All overlap, union and Venn analyses were performed at the Ensembl gene-ID level (version suffixes removed) to avoid symbol ambiguities. Gene-set enrichment was assessed by Fisher’s exact tests and ranked GSEA against MSigDB collections; multiple testing correction was applied (FDR). Pathway enrichment analysis was done using KEGG. YAP1/TAZ target genes were derived from the CHEA Transcription Factor Targets dataset via Harmonizome 3.0 ^58^. Detailed workflow for WES (CNV and somatic variants) data analysis is written in the Suppl. Methods.

### Statistical Analysis

All statistical analyses were conducted using GraphPad Prism 10 (GraphPad software Inc.). Statistical differences between 2 populations were calculated by unpaired t test (2-tailed). For multiple populations’ comparison, one-way or two-way ANOVA with Turkey’s multiple-comparisons test was used. p < 0.05 was considered statistically significant. 2-way inhibitor synergies were calculated using SynergyFinder 3.0 ^56^.

## Data Availability

The data that support the findings of this study are included with the manuscript and supplemental data files and are also available from the corresponding author upon reasonable request. RNA-Seq data from human meningioma cells expressing treated with either DMSO, IAG-933, or MYF-03-176 can be accessed at the GEO database at GSEXXXXXX.

## Acknowledgments

We thank Anna Spangler-Temby for continued administrative assistance. We thank Dr. Martin McMahon for insightful discussions. Funding for this study was provided by the University of Utah Department of Neurosurgery and Huntsman Cancer Institute Start-up funds (M.D., F.S.). We acknowledge support from Zaily Connell and Brian Dalley from the High-Throughput Genomics and HCI’s National Cancer Institute Cancer Center Support Grant P30CA042014. We would like to thank Tony Cooke and Kevin Jan, Yokogawa Corporation of America, for technical support.

## Author contributions

R.J., M.D., and F.S. conceived the study. D.K., M.B., D.S., E.I., S.M.J., A.G.P., D.R.V., M.F., and D.G. performed the experiments. D.K., M.B., D.S., S.M.J., Q.L., M.K.S., and F.S. analyzed the data. F.S., D.S., and S.H. wrote the manuscript. Q.L. and F.S. reviewed and edited the manuscript. W.C., R.J., M.D., and F.S. acquired the funding. D.N., R.D.R., H.C., W.C., R.J., and F.S. supervised the study. All authors read, reviewed, and approved the manuscript.

## Supplementary Tables

Suppl. Table S1: Enrichment of YAP1/TAZ-TEAD target genes in the differentially regulated genes identified in the six meningioma cell lines upon IAG-933 or MYF-03-176 treatment by RNA-Seq. Differentially expressed genes observed upon TEADi treatment (IAG-933, MYF-03-176) compared to DMSO treatment in BenMen1, BrMen, KT21-MG1, CH1-57MN, and IOMM-Lee cells.

Suppl. Table S2: Differentially expressed genes observed upon combined Y+T knockout compared to CD8A KO in BenMen1 and KT21-MG1 cells.

Suppl. Table S3: Significantly enriched KEGG terms in the six meningioma cell lines upon IAG-933 treatment.

Suppl. Table S4: CNV (A) and SV (B) data for cell lines with WES data.

Suppl. Table S5: Supplementary methods. A) Antibodies used B) sgRNA sequences C) Primers.

## Notes

### Competing Interest Statement

The authors have declared no competing interest.

### Summary of Updates

Figure 6 revised, added Figure 7. Updated author list

