## Supplementary material for "Comprehensive analysis of TEAD inhibition in meningioma identifies MEK and mTOR inhibition as effective combination therapies against resistant lines": Suppl. Figures

Suppl. Figures S1-S7

Suppl. Tables S1-S4

### Suppl. Figure S1

#### A Sanger sequencing of CRISPR-Cas9-edited CD8A, YAP1, and WWTR1 genes

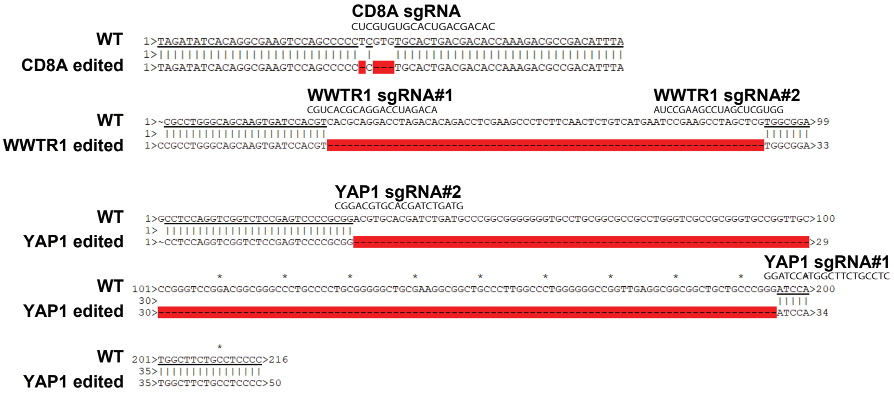


#### B Expression of YAP1/TAZ-TEAD target genes upon CRISPR-Cas9-mediated gene editing

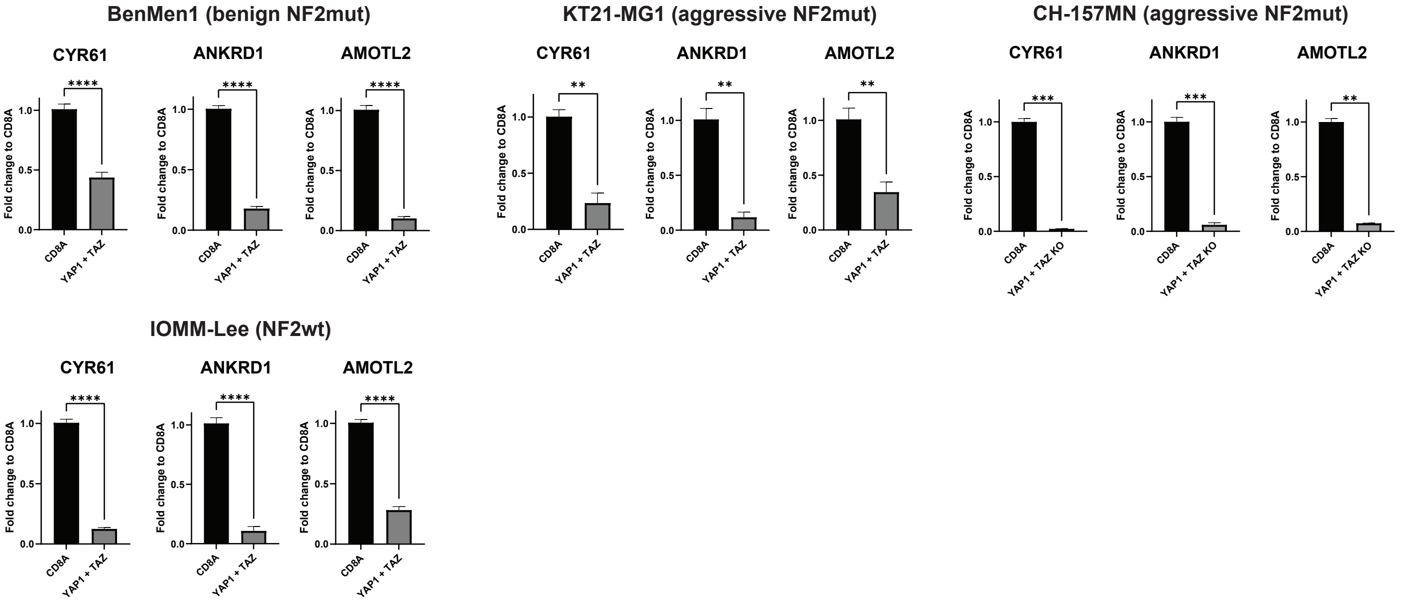

### Suppl. Figure S2 A-B

#### A Western blot showing NF2/Merlin protein expression in different meningioma cell lines

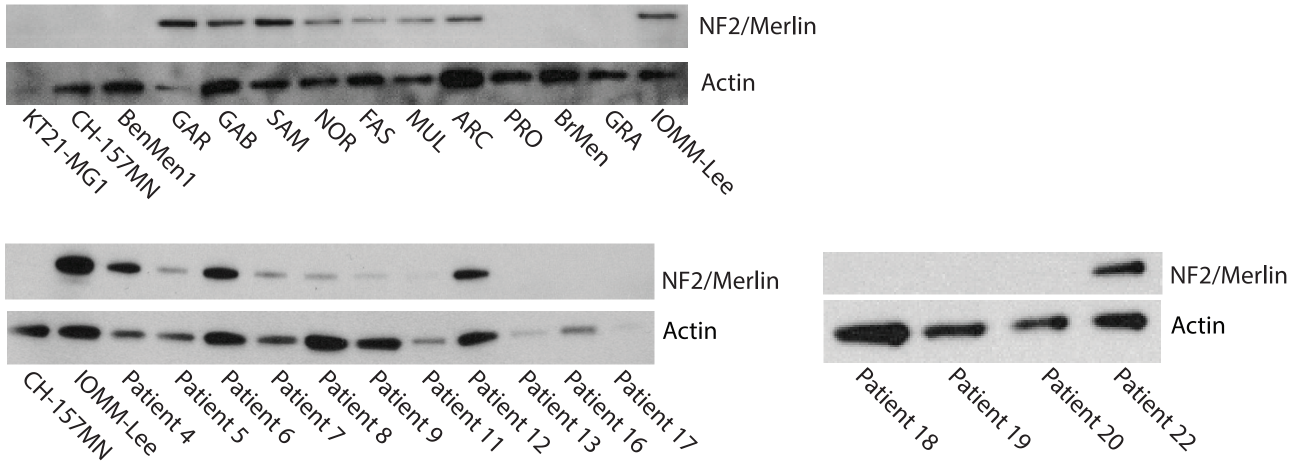


#### B Dose response curves for meningioma cell lines treated with four different TEAD inhibitors

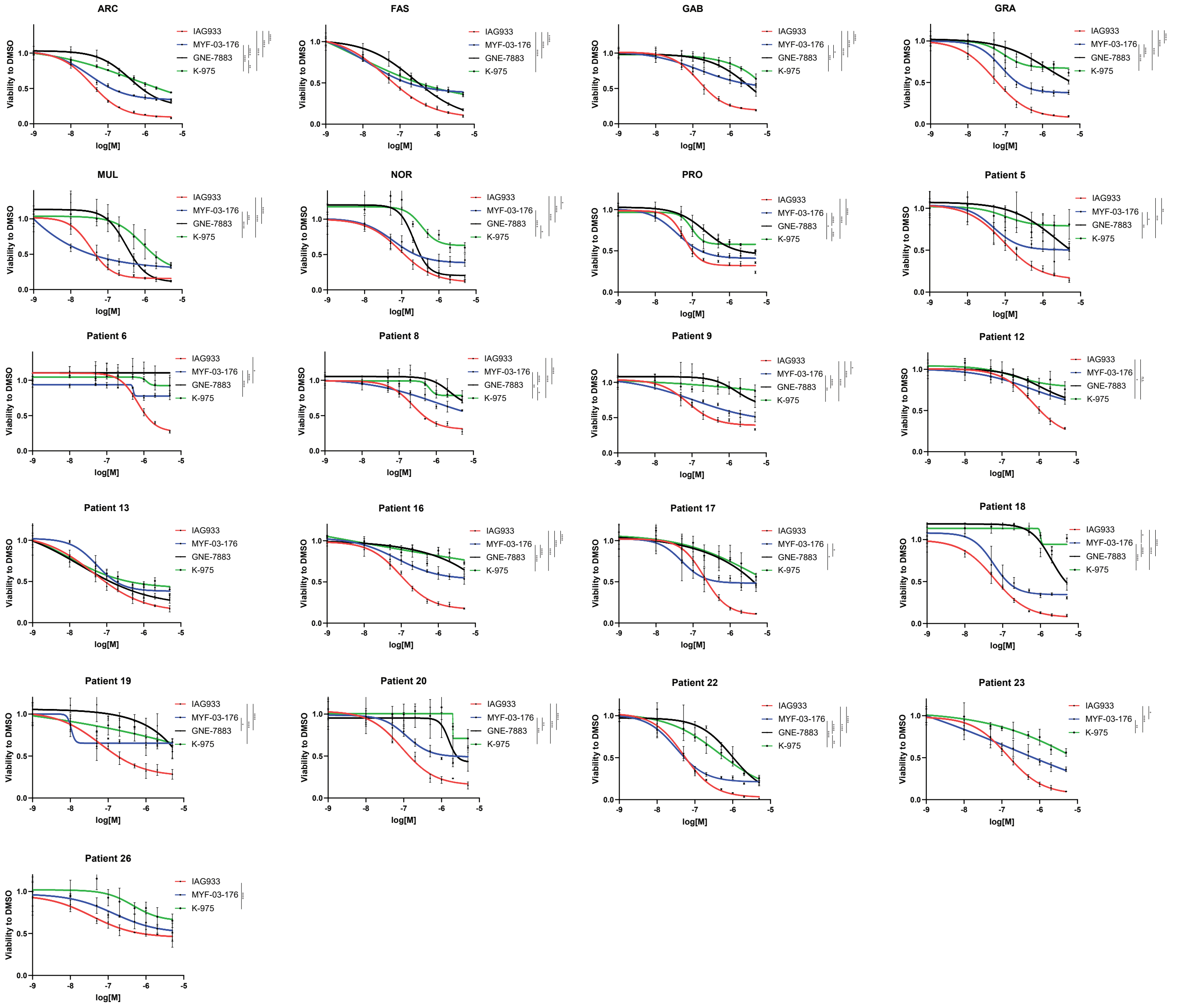

### Suppl. Figure S2 C-D

#### C Growth curves of meningioma cell lines treated with the TEAD inhibitor IAG-933

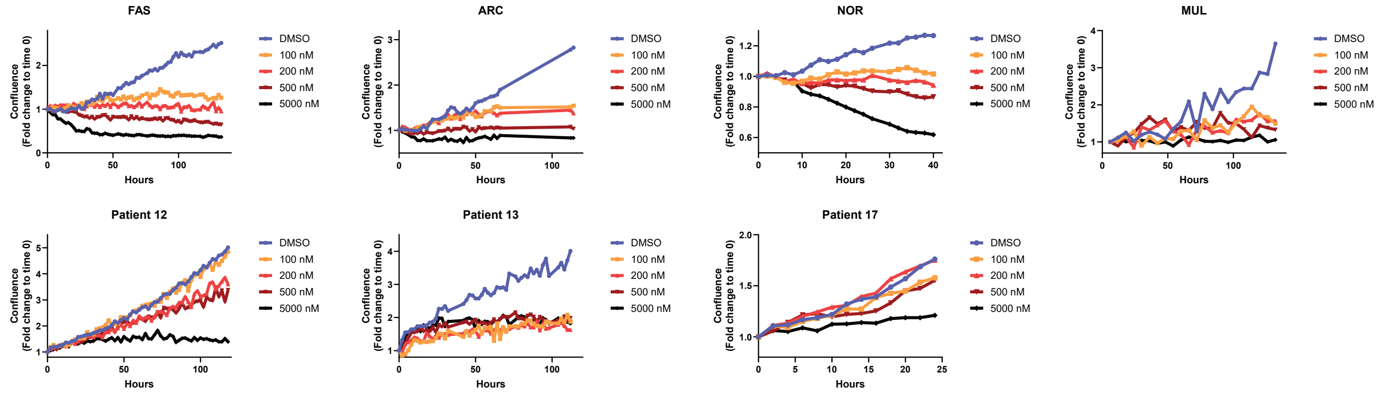


#### D Growth curves of meningioma cell lines treated with the TEAD inhibitor MYF-03-176

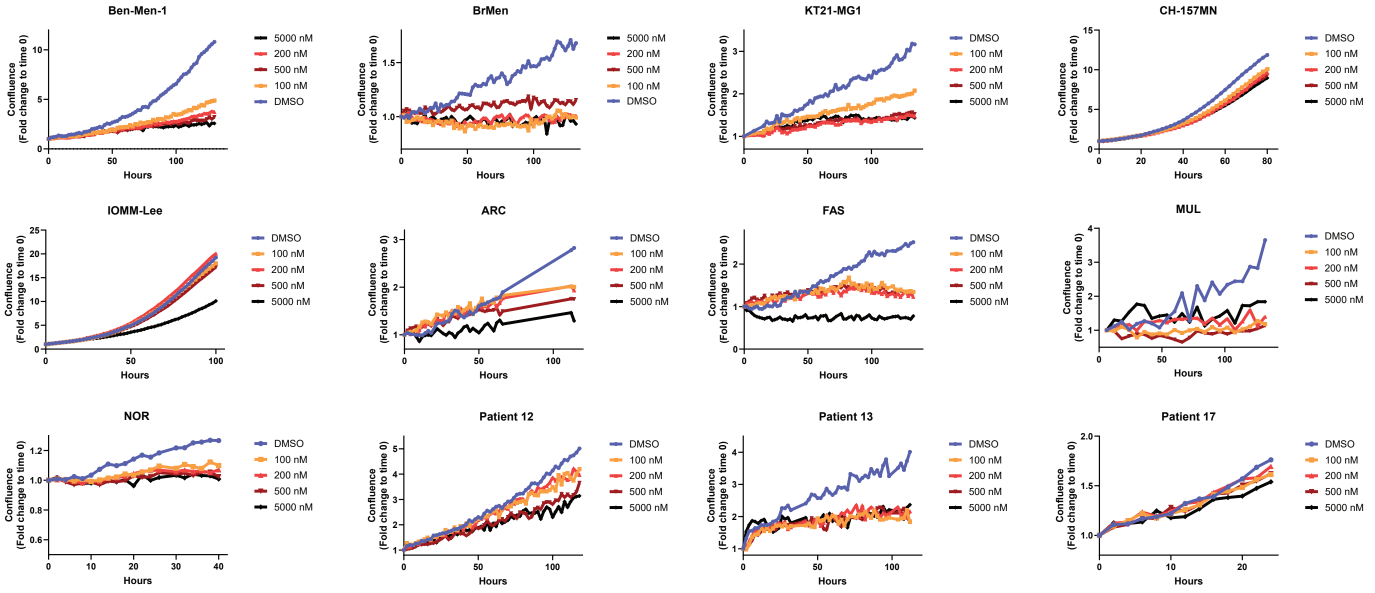

### Suppl. Figure S2 E-F

#### E Growth curves of meningioma cell lines treated with the TEAD inhibitor GNE-7883

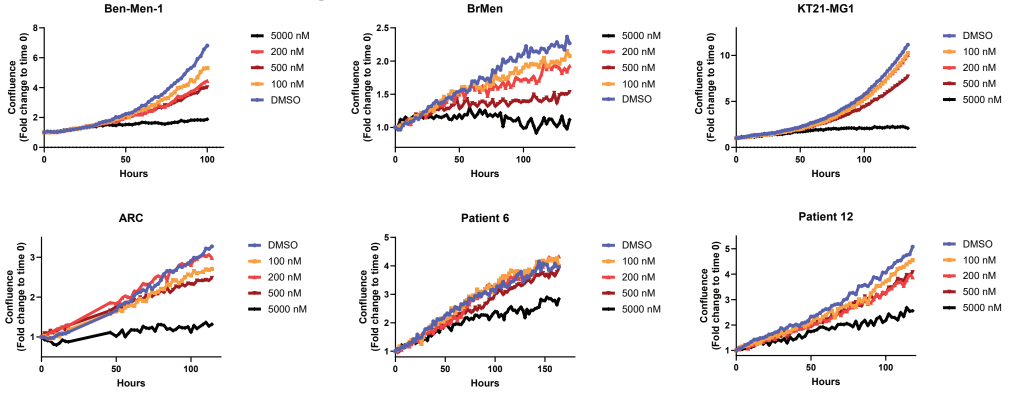


#### F Growth curves of meningioma cell lines treated with the TEAD inhibitor K-975

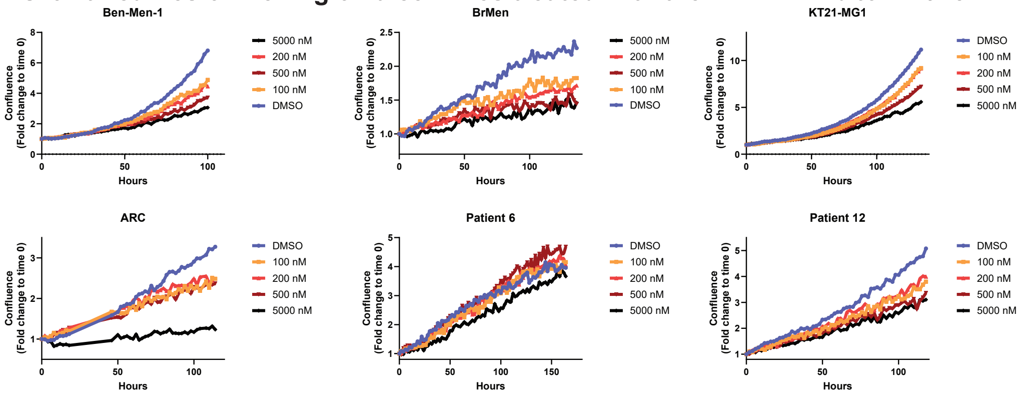

### Suppl. Figure S2 G

#### G Expression of YAP1/TAZ-TEAD target genes in meningioma cell lines upon treatment with four different TEAD inhibitors

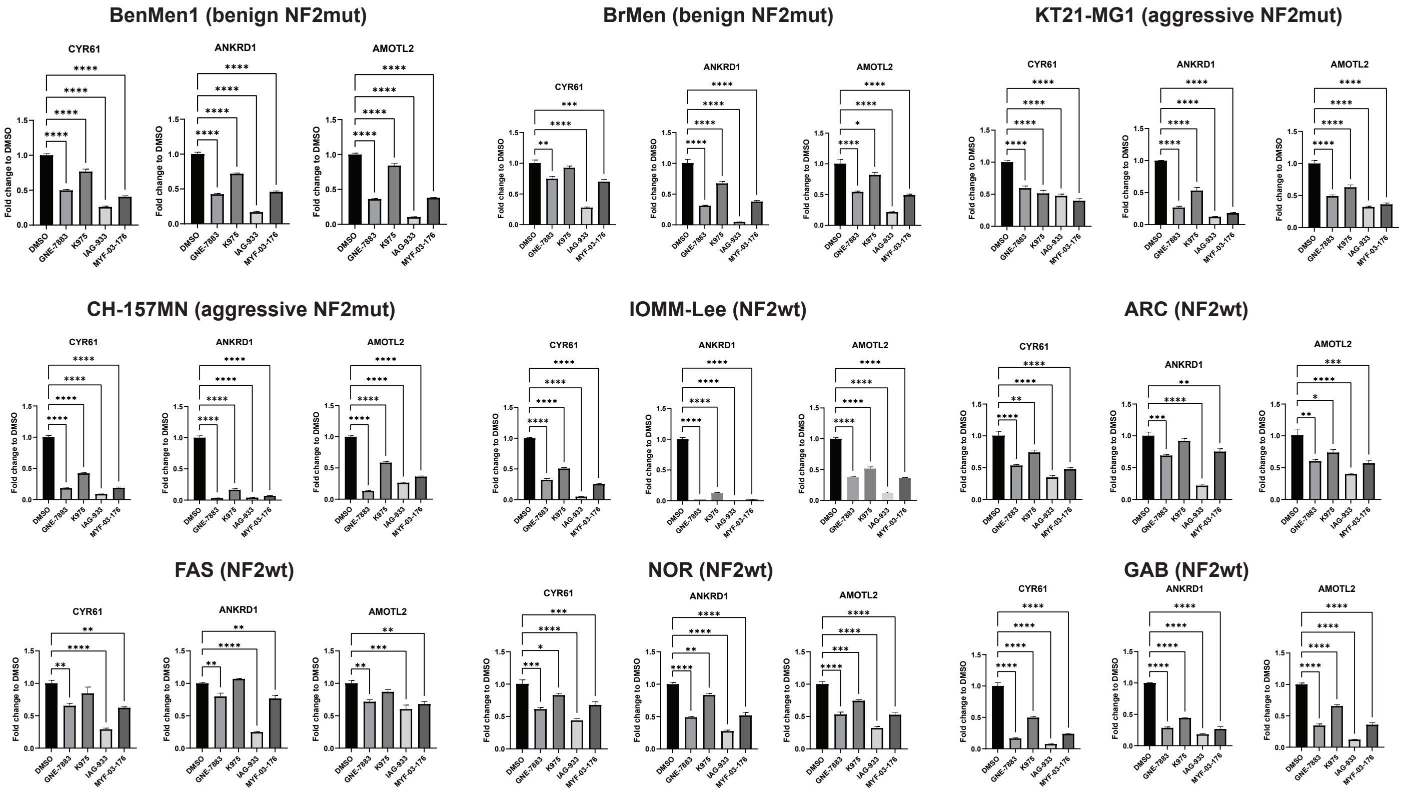

#### Suppl. Figure S3 A-C

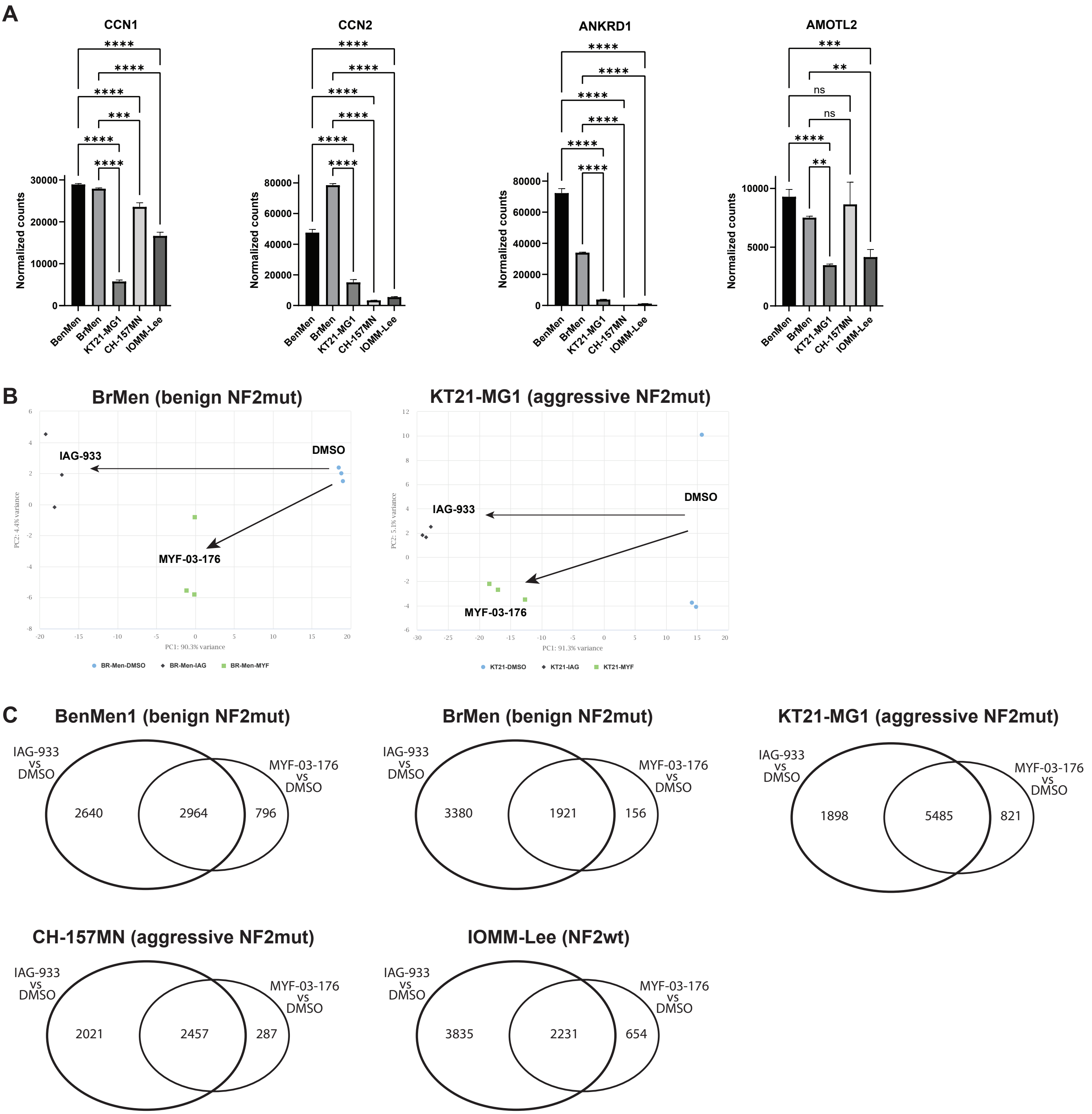

### Suppl. Figure S3 D

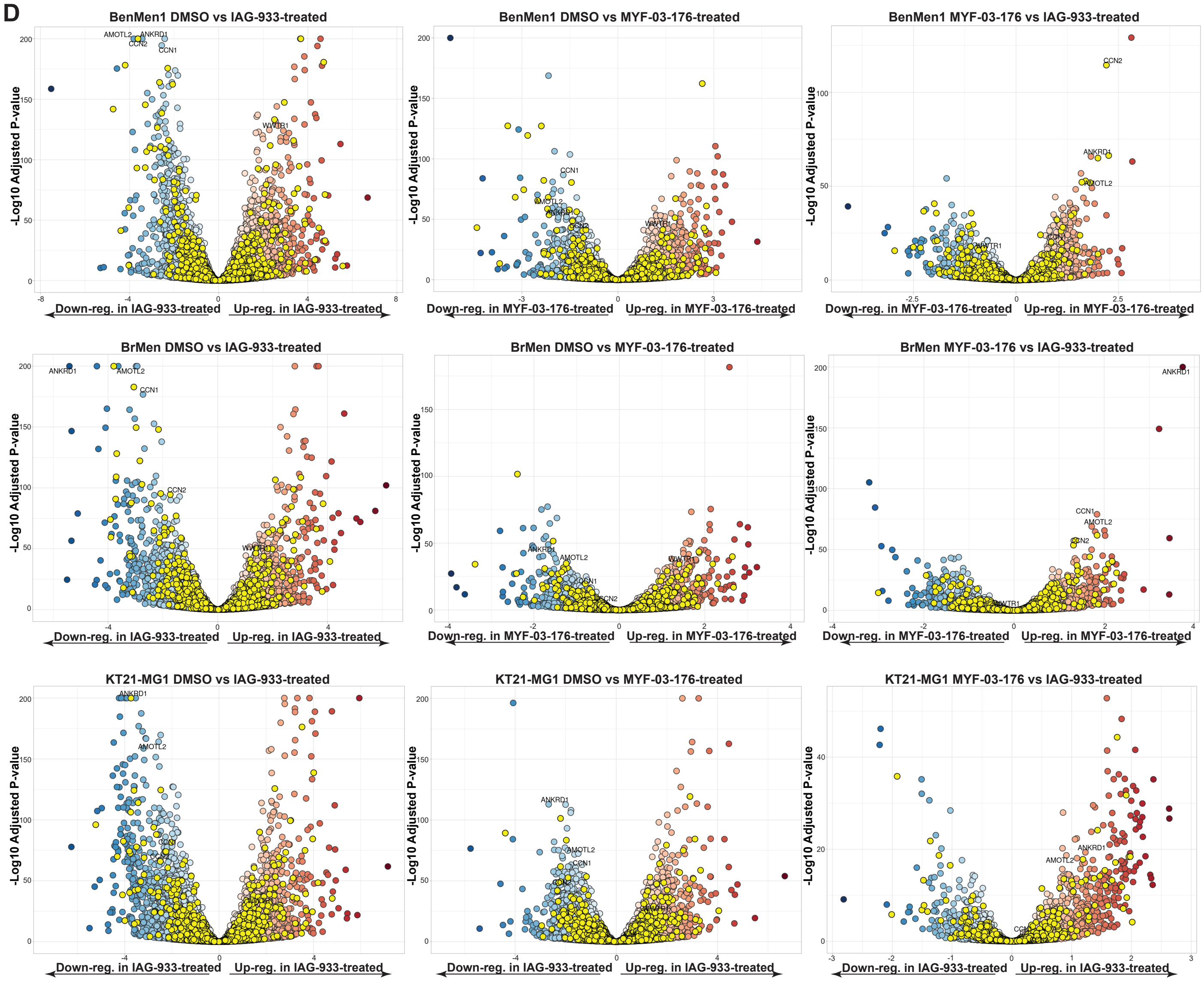

### Suppl. Figure S3 E-F

E

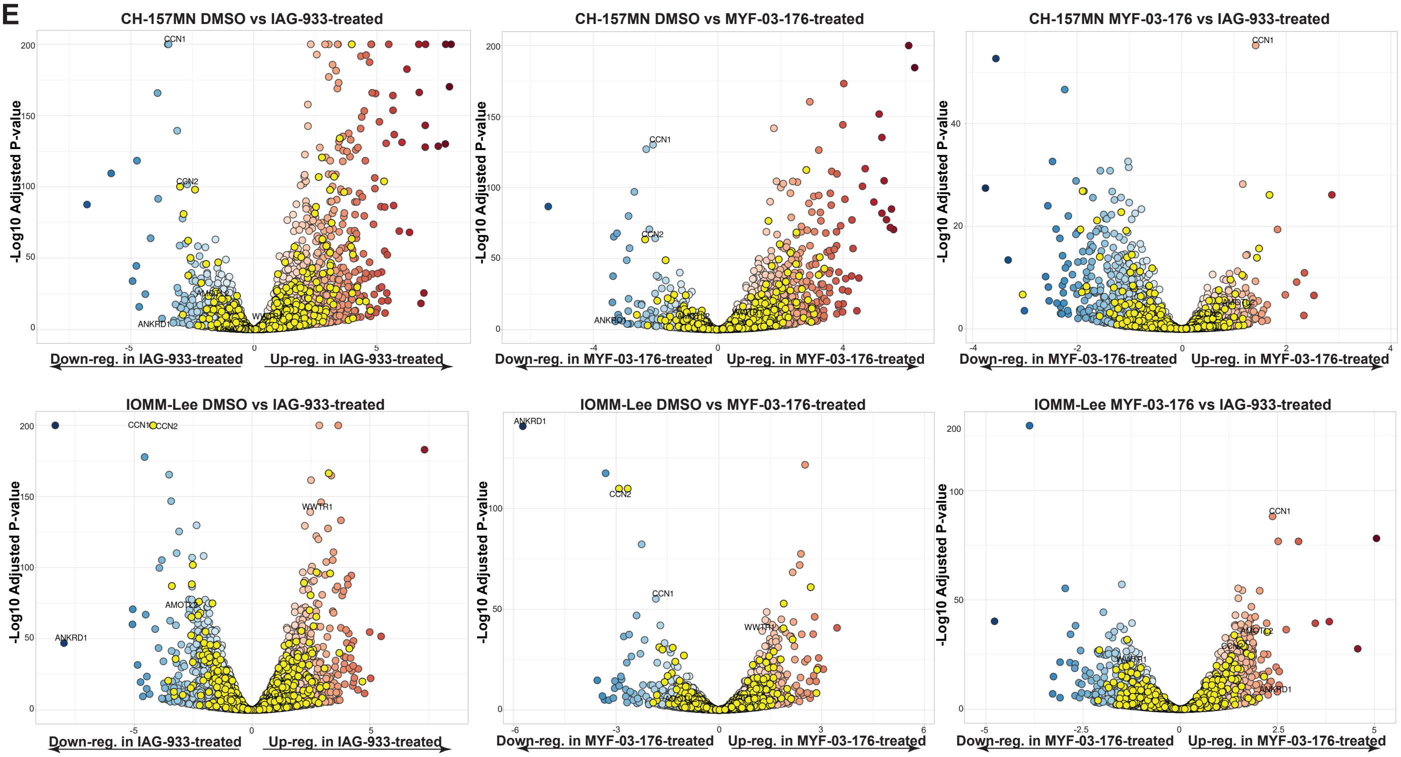


F

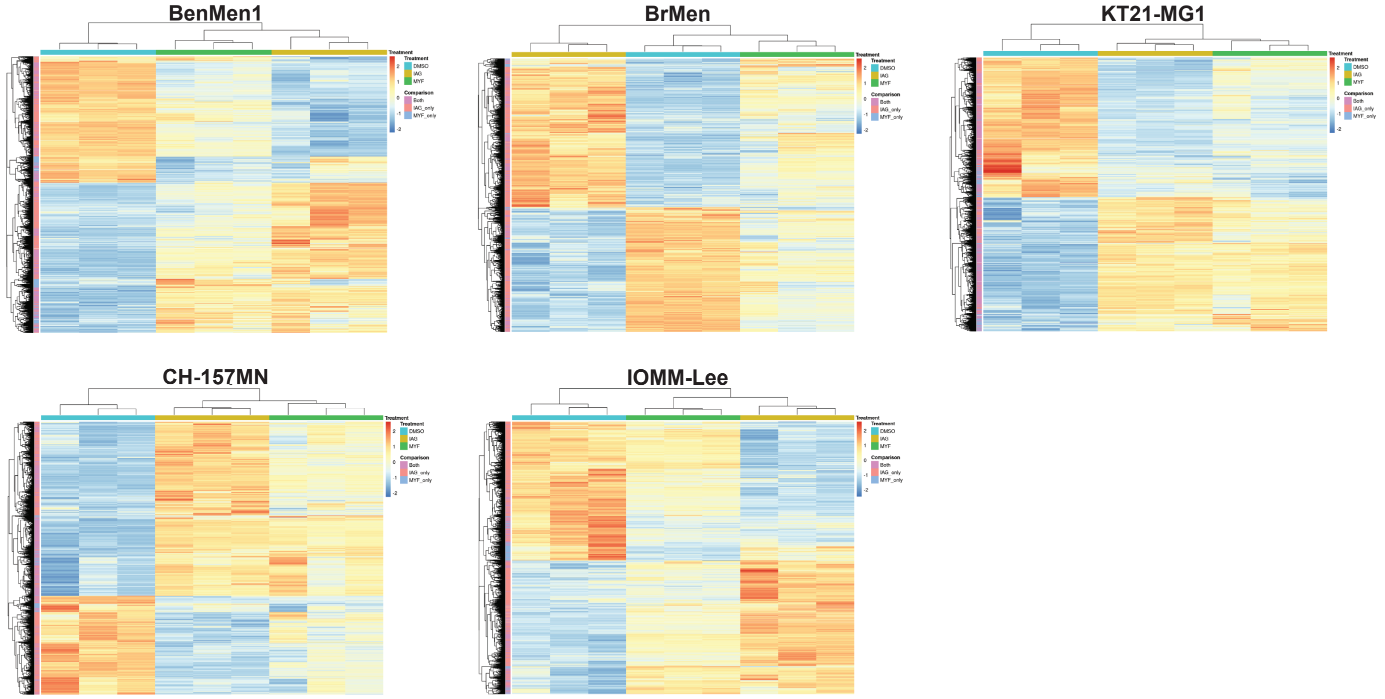

#### Suppl. Figure S4

##### A PCA of RNA-Seq data - CD8A knockout versus YAP1+TAZ double knockout

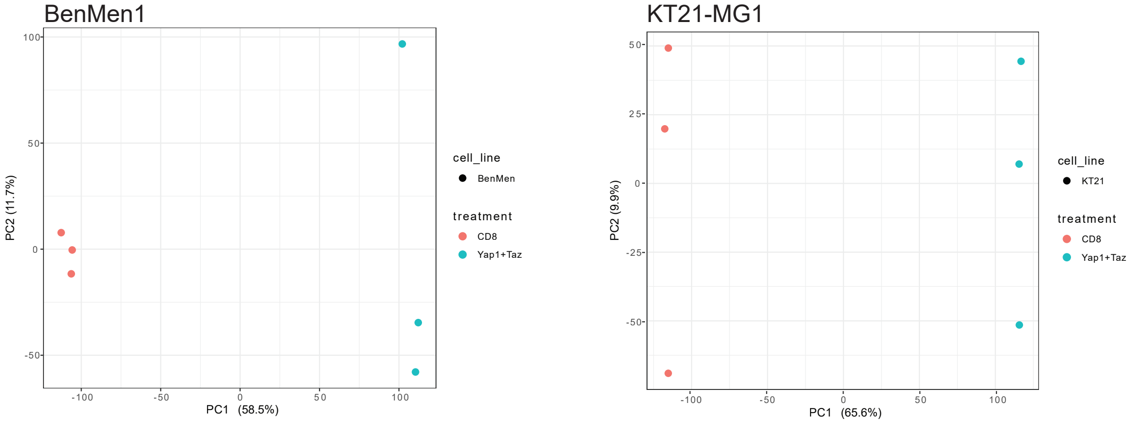


##### B Absolute expression of YAP1/TAZ target genes in BenMen1 cells upon TEADi or CRISPR-Cas9 editing

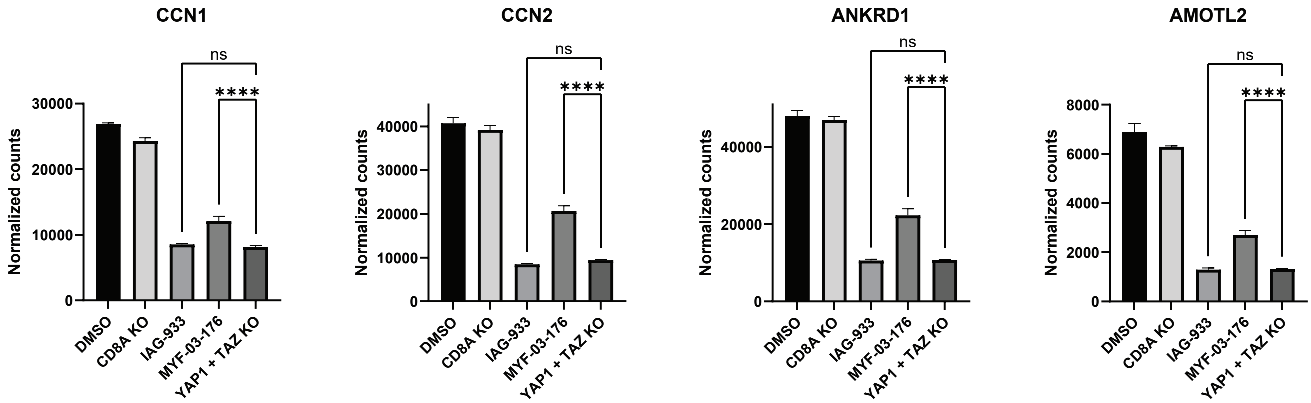


##### C Absolute expression of YAP1/TAZ target genes in KT21-MG1 cells upon TEADi or CRISPR-Cas9 editing

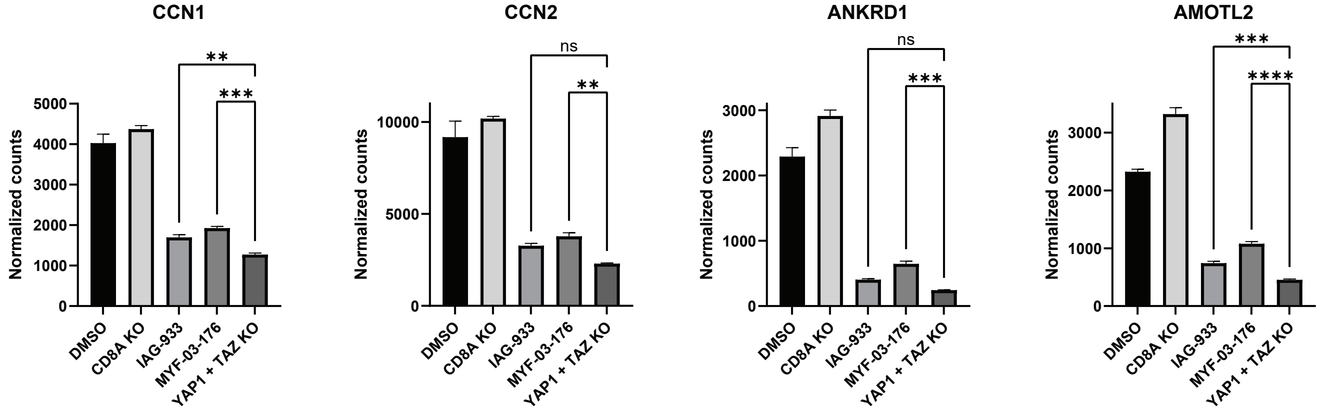


##### D Overlap DEGs TEADi vs YAP1/TAZ KO

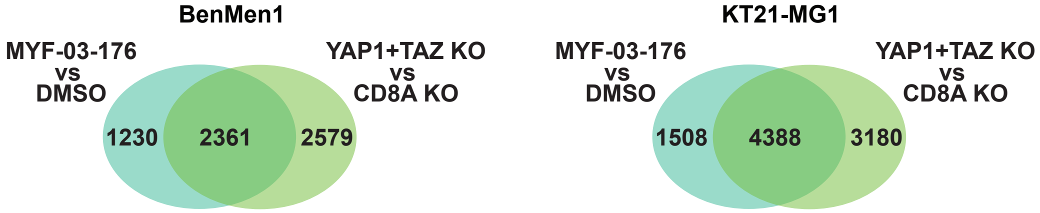

Suppl. Figure S5

**A** Activation of MAPK, PI3K-AKT-mTOR pathways in sensitive and resistant lines upon TEADi treatment

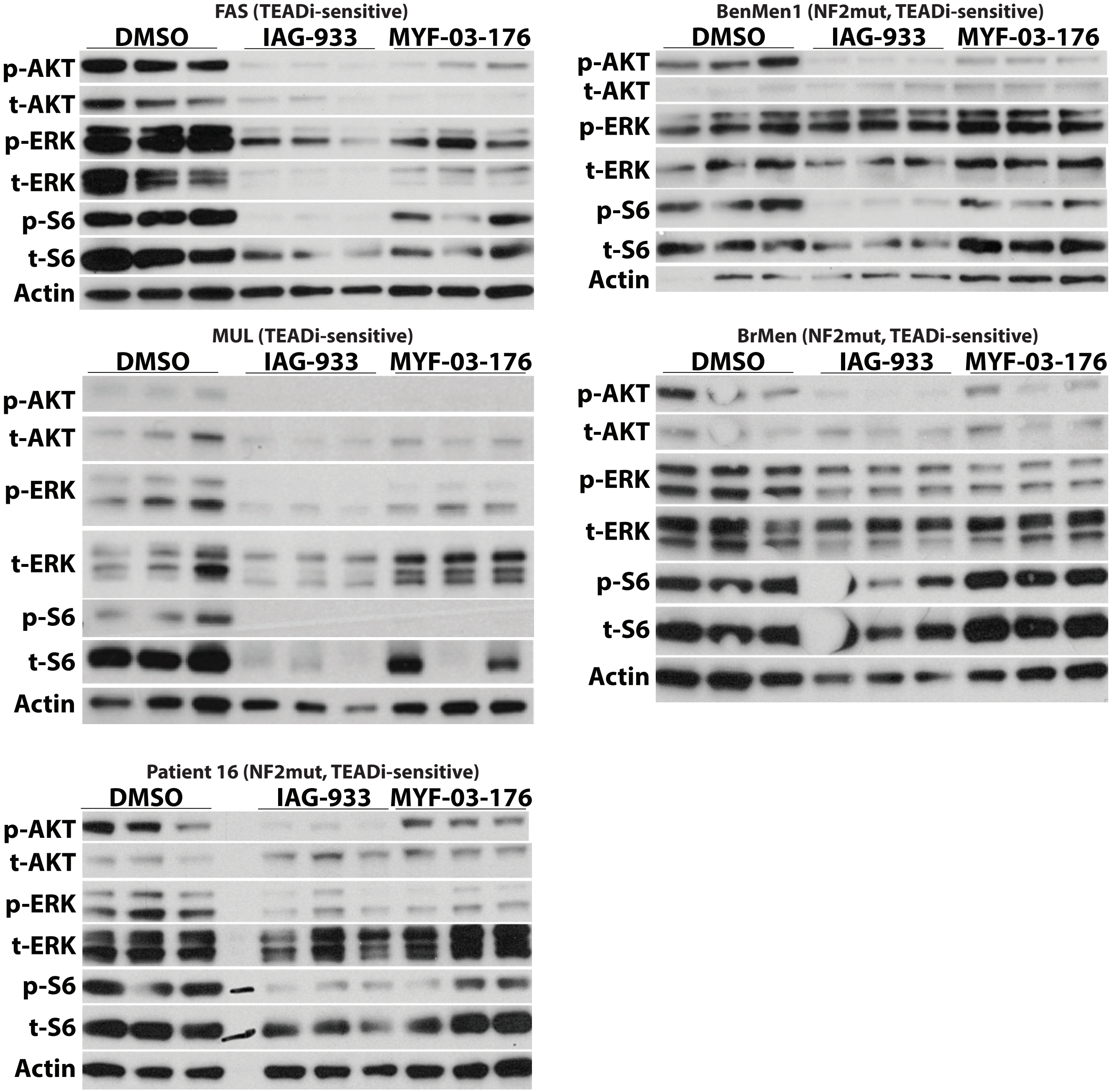

### Suppl. Figure S6 A-D

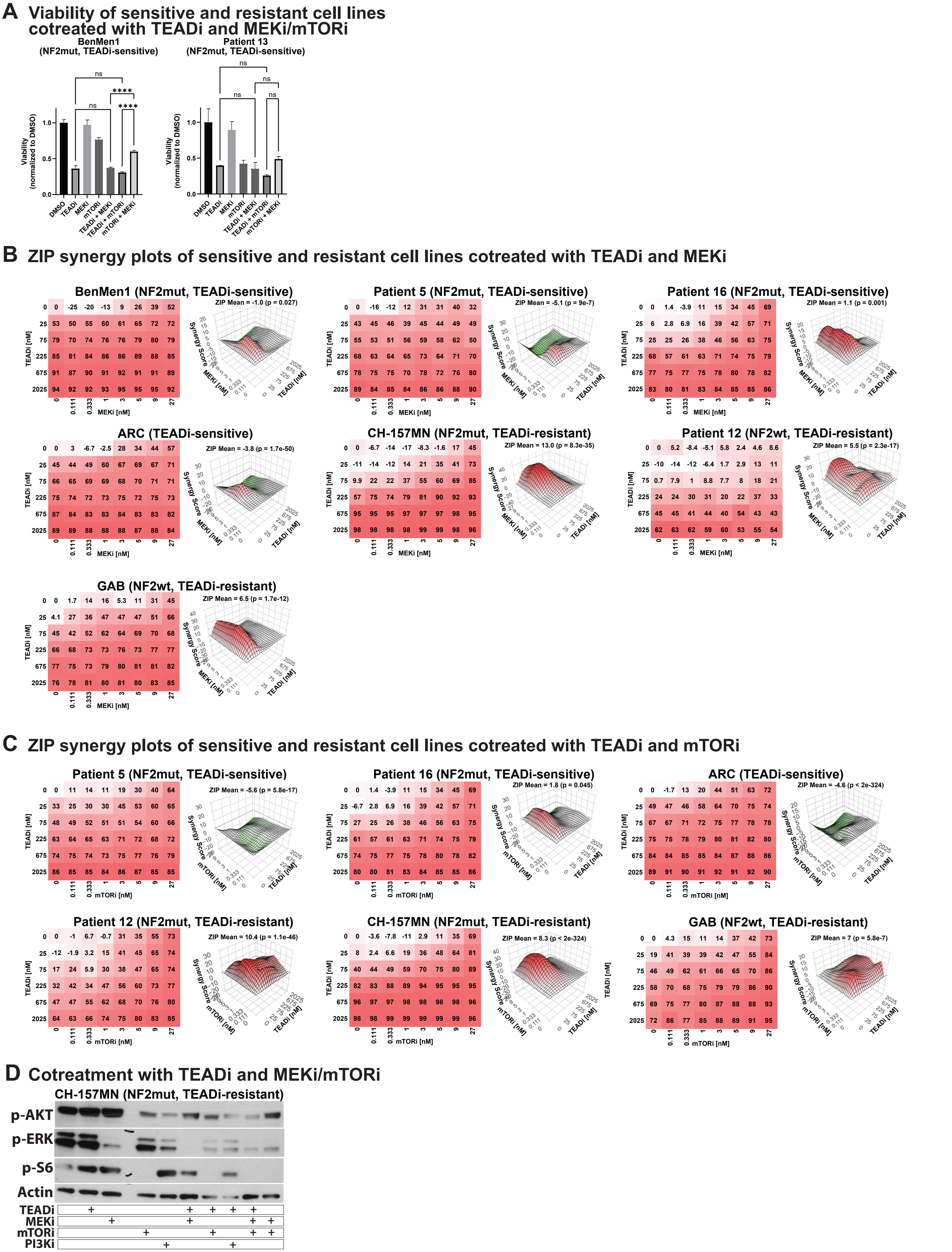

### Suppl. Figure S6 E-G

#### E Cotreatment with TEADi and FDA-approved mTORi

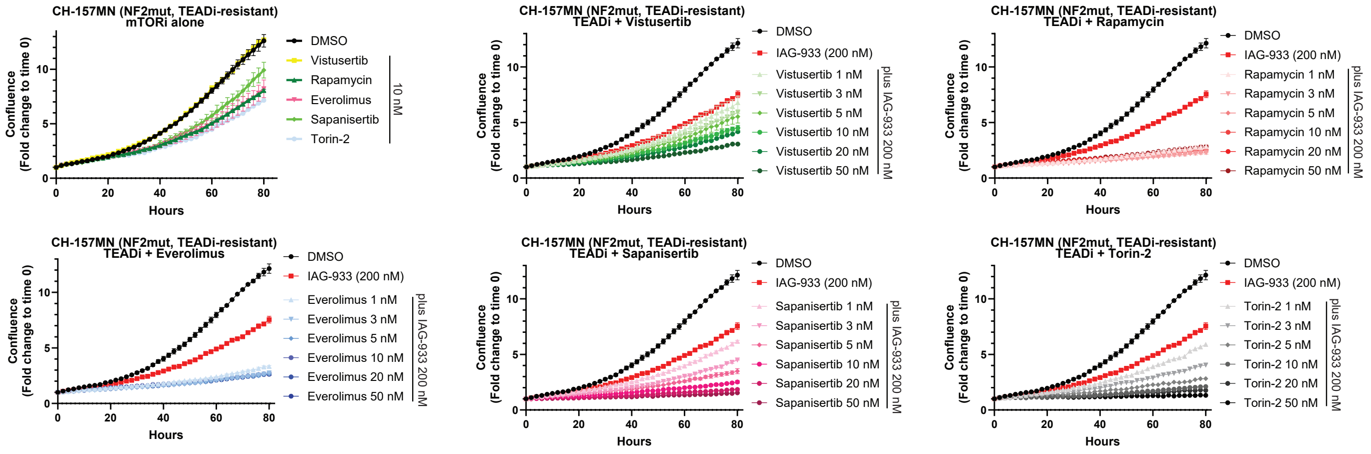


#### F Dose-response curves of meningioma cell lines treated with mTOR inhibitors and TEADi

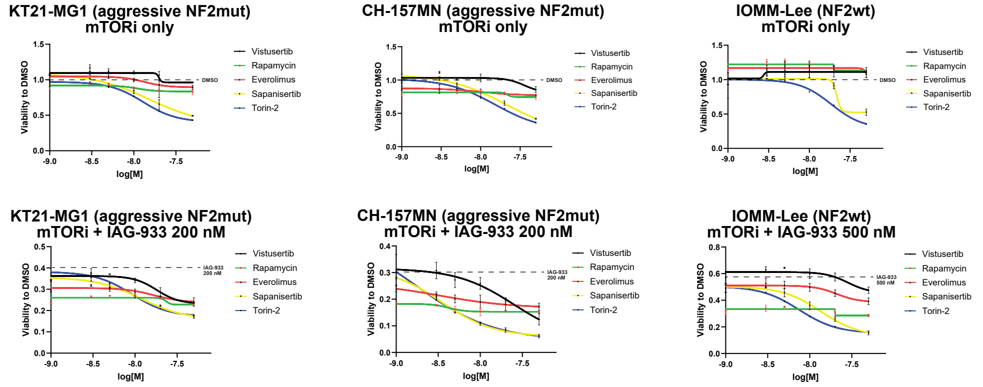


#### G Viability of cell lines cotreated with TEADi and FAKi

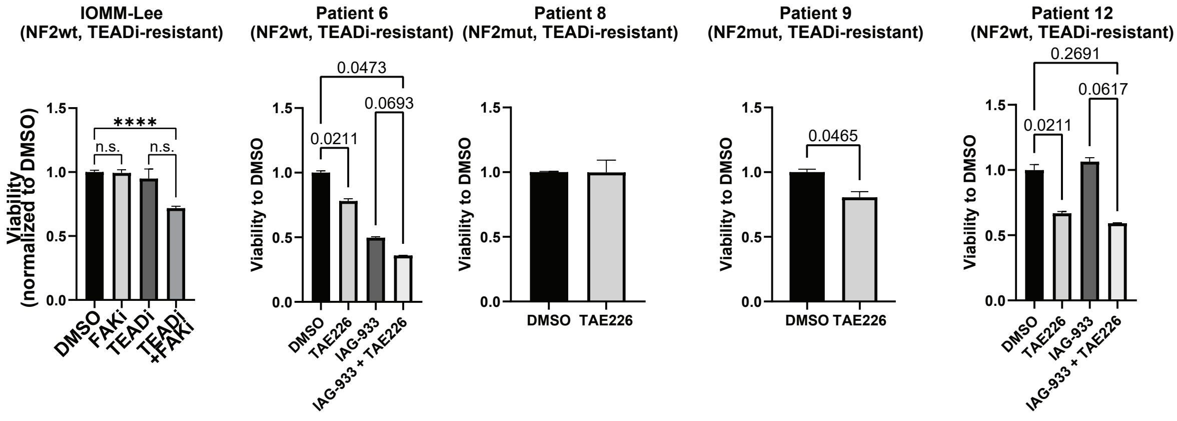

**Suppl. Figure S7**

**A**

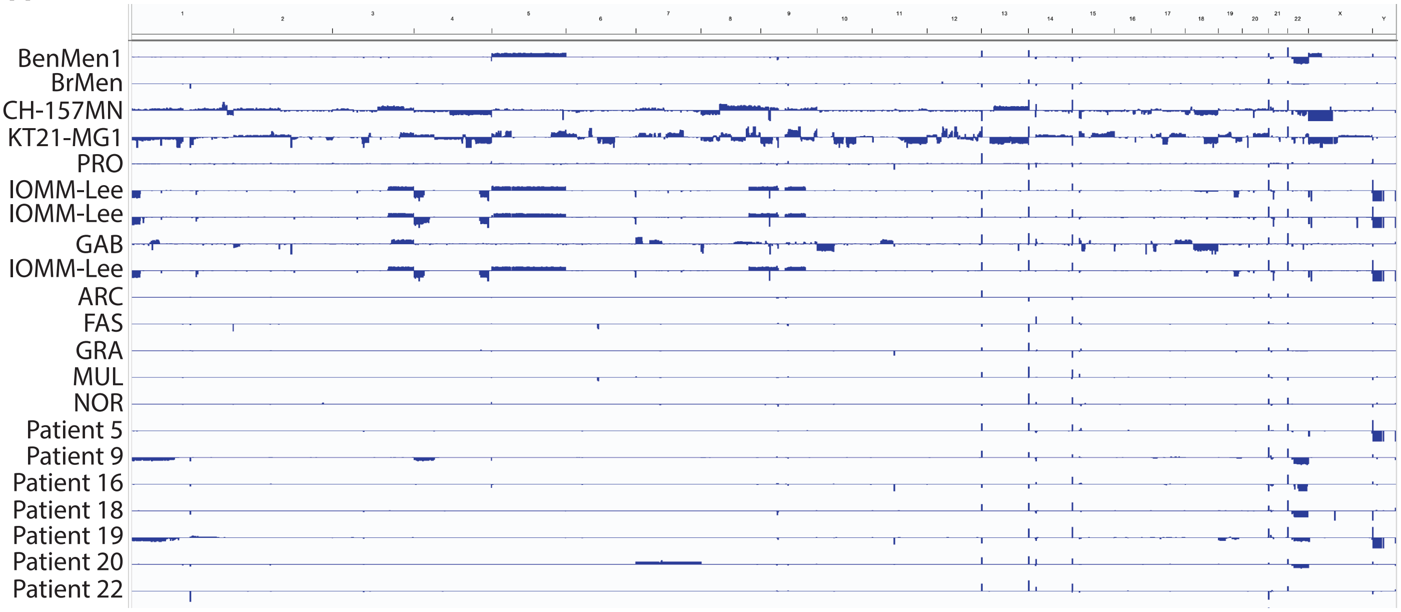

Suppl. Figure S1: Combined loss of YAP1 and TAZ (WWTR1) significantly reduces the growth of both NF2 mutant and NF2 wild type meningioma cells. A) Expression of the YAP1/TAZ-TEAD target genes CYR61, ANKRD1, and AMOTL2 in benign NF2mut (BenMen1 (n = 10 each)), aggressive high-grade NF2mut (KT21-MG1 (n = 3 each), CH-157MN (n = 3 each)), and NF2wt (IOMM-Lee (n = 14 each)) meningioma cell lines upon CRISPR-Cas9-mediated inactivation of either CD8A (control) or combined inactivation of YAP1 and TAZ (WWTR1). B) gDNA sequences of WT and edited (CD8A, YAP1, WWTR1/TAZ) gDNA samples. Error bars show SEM. Analysis was done using a two-tailed t test (A).

Suppl. Figure S2: TEADi shows significant efficacy against benign and a subset of high-grade NF2 mutant meningioma cell lines. A) Western Blot showing NF2 status in the meningioma cell lines used in this study. B) Viability dose response curves of meningioma cell lines treated with different doses of four TEAD inhibitors (IAG-933, MYF-03-176, GNE-7883, K-975). All experiments were performed in triplicates. C) Growth curves measured by live-cell imaging of meningioma cell lines treated with different doses of the TEAD inhibitor IAG-933 (100, 200, 500, 1000 nM, 0.05% DMSO). All experiments were performed in triplicates. D) Growth curves measured by live-cell imaging of meningioma cell lines treated with different doses of the TEAD inhibitor MYF-03-176 (100, 200, 500, 1000 nM, 0.05% DMSO). All experiments were performed in triplicates. E) Growth curves measured by live-cell imaging of meningioma cell lines treated with different doses of the TEAD inhibitor GNE-7883 (100, 200, 500, 1000 nM, 0.05% DMSO). All experiments were performed in triplicates. F) Growth curves measured by live-cell imaging of meningioma cell lines treated with different doses of the TEAD inhibitor K-975 (100, 200, 500, 1000 nM, 0.05% DMSO). All experiments were performed in triplicates. G) Expression of the YAP1/TAZ-TEAD target genes CYR61, ANKRD1, AMOTL2 in meningioma cell lines upon treatment with four TEAD inhibitors (IAG-933 (500 nM), MYF-03-176 (500 nM), GNE-7883 (5000 nM), K-975 (1000 nM)). All experiments were performed in triplicates. Error bars show SD (A) or SEM (F). Analysis was done using an ordinary one-way ANOVA (F) or ordinary two-way ANOVA (A, comparison at 200 nM) with multiple comparisons testing.

Suppl. Figure S3: TEADi treatment results in a downregulation of YAP1/TAZ-TEAD signaling in both TEADi-sensitive and -resistant cell lines. A) Baseline expression of YAP1/TAZ-TEAD target genes in five meningioma cell lines. B) PCA plots showing DMSO, IAG-933, or MYF-03-176-treated samples of benign NF2mut (BrMen, left) or high-grade NF2mut (KT21-MG1, right) meningioma cell lines. C) Venn diagram showing overlap between DEGs induced by IAG-933 or MYF-03-176 treatment (compared to DMSO) for the different cell lines. D-E) Volcano plot showing differentially expressed genes in TEADi-sensitive cell lines (BenMen1, BrMen, KT21-MG1) (D) and TEADi-resistant cell lines (CH-157MN, IOMM-Lee) (E) between samples treated with DMSO or TEADi (IAG-933, MYF-03-176). Yellow dots represent known YAP1/TAZ-TEAD target genes. F) Heatmaps showing expression of significantly regulated genes in DMSO, IAG-933, and MYF-03-176-treated samples in the six meningioma cell lines.

Suppl. Figure S4: TEADi induces highly similar transcriptional changes compared to YAP1/TAZ knockout. A) PCA plots showing CD8A KO and Y+T KO samples from BenMen1 and KT21-MG1 cell lines. B-C) Absolute expression of the YAP1/TAZ target genes CCN1, CCN2, ANKRD1, and AMOTL2 in DMSO/TEADi-treated and CD8A/Y+T KO samples from BenMen1 (B) and KT21-MG1 (C) cell lines. D) Overlap of DEGs from MYF-03-176-treated samples (compared to DMSO) and

Y+T KO samples (compared to CD8A KO) for BenMen1 and KT21-MG1 cell lines. All experiments were performed in triplicates. Error bars show SD (B-C). Analysis was done using an ordinary one-way ANOVA (B-C) with multiple comparisons testing.

Suppl. Figure S5: TEADi induces MEK-ERK and mTOR-S6 (but not PI3K-AKT) signaling in TEADi-resistant cell lines. A) Representative western blots showing phosphorylated and total protein levels of AKT, ERK, and S6 in TEADi-sensitive meningioma cell lines upon treatment with either DMSO (control) or TEADi (IAG-933 or MYF-03-176, 500 nM each). Actin was used as a loading control.

Suppl. Figure S6: Co-targeting MAPK, mTOR-S6, or FAK signaling is able to overcome resistance to TEADi. A) End point viability of TEADi-sensitive meningioma cell lines upon co-treatment with combinations of TEADi (IAG-933, 200 nM), PI3Ki (pictilisib, 50 nM), MEKi (trametinib), or mTORi (torin-2). All experiments were performed in triplicates. B-C) Synergy plots for TEADi/MEKi (B) or TEADi/mTORi (C) combination treatments of TEADi-sensitive or -resistant meningioma cell lines. D) Western Blot showing activation of MAPK (p-ERK), PI3K-AKT (p-AKT), or mTOR-S6 (p-S6) pathways in TEADi-resistant CH-157MN cells upon treatment with TEADi (IAG-933), MEKi (trametinib), mTORi (torin-2), or PI3Ki (pictilisib) in various combinations. Actin was used as a loading control. E) Growth curves of TEADi-resistant CH-157MN cells upon treatment with mTOR inhibitors alone (10 nM each) or combination treatment of TEADi (IAG-933, 200 nM) plus additional mTOR inhibitors (torin-2, rapamycin, vistusertib, everolimus, sapanisertib, dose range of 1-50 nM). All experiments were performed in duplicates. F) Dose-response curves of meningioma cell lines (KT21-MG1, CH-157MN, IOMM-Lee) treated with mTOR inhibitors (see F) alone or in combination with TEADi (IAG-933, 200 nM or 500 nM). All experiments were performed in duplicates. G) End point viability of TEADi-sensitive and -resistant cell lines upon treatment with TEADi (IAG-933, 200 nM) or FAKi (TAE226, 500 nM). Error bars show SEM (A, E, F). Analysis was done using an ordinary one-way ANOVA (A,G) or ordinary two-way ANOVA (F (comparison at last time point)) with multiple comparisons testing. Synergy analysis for two-way treatments was done with SynergyFinder3.0.

Suppl. Figure S7: Figure 7: Whole exome sequencing identifies Ras-Raf pathway mutations in TEADi resistant cell lines. A) Manhattan plot showing CNVs in the 19 meningioma cell lines with WES data.

Supplementary methods:

Code Availability

Analysis code for RNA-Seq analysis is available at https://github.com/qingl0331/A8783-27033R-vs-26519R. Analysis code for WES analysis is available under https://github.com/HuntsmanCancerInstitute/Workflows/tree/master/Hg38RunnerWorkflows2/DnaAlignQC/AutoAnalysis and https://github.com/HuntsmanCancerInstitute/Workflows/tree/master/Hg38RunnerWorkflows2/SomaticCaller and https://github.com/HuntsmanCancerInstitute/Workflows/tree/master/Hg38RunnerWorkflows2/CopyAnalysis/TumorOnly.

Bioinformatic Analysis of whole exome sequencing data

Bioinformatic analysis of the Meningioma project samples made use of a series of Apptainer containerized Snakemake workflows to process each tumor cell line's low depth whole genome sequence (WGS), high depth exome sequencing, and bulk RNASeq datasets. These are publicly available from the Huntsman Cancer Institute GitHub repository and are optimized to run on a Slurm cluster in the HIPAA compliant protected environment at the Center for High Performance computing (https://chpc.utah.edu).

The DnaAlignQC workflow takes exome or WGS paired fastq sequence reads, removes adapter sequence (Trimmomatic), aligns each read pair to GRCh38 (bwa-mem2, GRCh38_GIABv3_no_alt_analysis_set_masked), filters the alignments for quality, unique mapability, and removes duplicates (Samtools fixmate, view, markdup). DnaAlignQC also calculates a variant of quality control metrics including, uniformity to accurately estimate the alignment read depth (USeq UniObRC), cross species contamination (fastq_screen), and sequencing quality (fastqc). Lastly, QC metrics are aggregated across all samples into reports and spreadsheets (MultiQC, USeq AggregateQCStats2).

To call somatic SNV and INDEL variants from the tumor exome samples, the SomaticCaller workflow was utilized. This first runs Illumina's Manta and Strelka callers in somatic mode producing variants that are subsequently filtered for variant quality (tumor AF>= 0.05, DP>=20, UniOb>=3; BKZ >=4, USeq/StrelkaFilter). BKZ is a variant specific, z-score background error model (USeq VCFBkz) used to weight a variant's QUAL score based on an internally generated set of normal exome samples. Variants passing the quality filters were annotated with SnpEff, CLINVAR, gnomAD, and detailed splice site info (USeq VCFSpliceScanner) using the SpliceAI, OncoKB, and Annotator workflows. Variants with a gnomAD population allele frequency of > 0.01 or those with labeled as benign or likely_benign by CLINVAR or likely_neutral by OncoKB where excluded. Variants must also pass one or more of the following criteria: being labeled by SnpEff as a HIGH impact - loss of function mutation, marked as pathogenic or likely-pathogenic by CLINVAR, or oncogenic, likely_oncogenic, or resistance by OncoKB, or having a MaxEntScan or SpliceAI damaged splice junction. These methods generate high quality and high consequence somatic SNV and INDEL variants.

Somatic CNVs from the WGS datasets were identified using the tumor only CopyAnalysis workflow. This follows GATK's best practice protocol to compare copy ratios for each window against a sex matched panel of normals generated internally to model expected coverage depth and variance. The workflow was run three times for each tumor sample using different window sizes (10KB, 25KB, and 100KB), filtered for windows with a tumor copy ratio >= 0.15, and combine into a final annotated call set (USeq CopyAnalysisBedMerger). To generate the germline vcf input for GATK, each tumor's WGS and exome fastq were combine, run through the DnaAlignQC workflow, and germline variants called using the Illumina Germline workflow in batch mode with all samples. SNVs and INDELs were filtered for a read depth >=6, alternative observations >=3, variant allele fraction >= 0.35, and a gnomAD population allele fraction < 0.01.

Applications and tool packages:

| | |
|---|---|
| Apptainer | https://github.com/apptainer/apptainer |
| Snakemake | https://github.com/snakemake/snakemake |

Trimmomatic https://github.com/usadellab/trimmomatic
bwa-mem2 https://github.com/bwa-mem2/bwa-mem2
USeq https://github.com/HuntsmanCancerInstitute/USeq
fastq_screen https://github.com/StevenWingett/FastQ-Screen
fastqc https://github.com/s-andrews/fastqc
MultiQC https://github.com/multiqc/multiqc
Samtools https://github.com/samtools/samtools
GRCh38_GIABv3 https://www.nist.gov/programs-projects/genome-bottle
Manta https://github.com/Illumina/manta
Strelka https://github.com/Illumina/strelka
GATK Somatic Copy Ratio https://gatk.broadinstitute.org/hc/en-us/articles/360035531092--How-to-part-I-Sensitively-detect-copy-ratio-alterations-and-allelic-segments
SpliceAI https://github.com/Illumina/SpliceAI
STAR https://github.com/alexdobin/STAR
featureCounts https://subread.sourceforge.net/
Clumpify https://bbmap.org/tools/clumpify
Cutadapt https://github.com/marcelm/cutadapt/
Ensembl https://www.ensembl.org/
Picard https://github.com/broadinstitute/picard

Workflows:
DnaAlignQC
https://github.com/HuntsmanCancerInstitute/Workflows/tree/master/Hg38RunnerWorkflows2/DnaAlignQC/AutoAnalysis
SomaticCaller
https://github.com/HuntsmanCancerInstitute/Workflows/tree/master/Hg38RunnerWorkflows2/SomaticCaller
IlluminaGermline
https://github.com/HuntsmanCancerInstitute/Workflows/tree/master/Hg38RunnerWorkflows2/IlluminaGermline
CopyAnalysis PoN
https://github.com/HuntsmanCancerInstitute/Workflows/tree/master/Hg38RunnerWorkflows2/CopyAnalysis/PoN
CopyAnalysis TumorOnly
https://github.com/HuntsmanCancerInstitute/Workflows/tree/master/Hg38RunnerWorkflows2/CopyAnalysis/TumorOnly
Annotator
https://github.com/HuntsmanCancerInstitute/Workflows/tree/master/Hg38RunnerWorkflows2/Annotator
SpliceAI
https://github.com/HuntsmanCancerInstitute/Workflows/tree/master/Hg38RunnerWorkflows2/SpliceAI
RnaAlignQC
https://github.com/HuntsmanCancerInstitute/Workflows/tree/master/Hg38RunnerWorkflows2/RnaAlignQC/AutoAnalysis
